# Biosensor-guided genome-wide CRISPR screens enable *cis*,*cis*-muconic acid overproduction in yeast

**DOI:** 10.64898/2026.10.31.762157

**Authors:** Mohamed A. Nasr, Adrien Knoops, Mackenzie Thornbury, Meghan Davies, David H. Kwan, Vincent J.J. Martin

## Abstract

Classical pathway metabolic engineering relies on enhancing strain productivity through the introduction of rationally identified genetic perturbations, which while useful, are time-consuming and miss unintuitive targets. Genome-wide CRISPR screens offer an alternative by efficiently sampling genomes, enabling the engineering of novel phenotypes, and are further enhanced when combined with high-throughput strain interrogation methods. Here, using *Saccharomyces cerevisiae*, we engineered small molecule biosensors into population enrichment tools using zeocin resistance and fluorescence as signal outputs. We then introduced the multi-functional genome-wide CRISPR (MAGIC) library that targets every gene for activation, interference, or deletion, thus creating biosensor-aided MAGIC, or senMAGIC. As a case study, we utilized senMAGIC to identify single genetic determinants that improve productivity of the nylon precursors protocatechuic and *cis*,*cis*-muconic acids (PCA and ccM). These were then randomly assembled as triple gRNA-tRNA arrays for multiplexed gene perturbations to identify combinatorial effects. Top gRNA hits were then confirmed using direct genomic integrations in senMAGIC-free production strains, resulting in identifying known as well as novel targets that increase PCA and ccM titres. Our results demonstrate senMAGIC as a robust automation-friendly whole-genome evolution tool and highlight limitations of the method, providing a roadmap for future multifunctional CRISPR screens to optimize host metabolism for efficient bioprocesses.

## Introduction

Microbes have been successfully engineered and transformed into “cell factories” to produce natural and new-to-nature molecules (Nielsen, 2013; Van Dien, 2016). Yields from these engineered microbes have not always met market needs, a challenge that is especially important to allow low-value chemicals such as polymer precursors to compete with unsustainable low-cost petroleum-derived sources (Keasling, 2010; Y. Liu & Nielsen, 2019). Traditional metabolic pathway engineering methods to enhance strain performance usually involve the introduction of rationally identified modifications that modulate target genes resulting in an increase, reduction, or ablation of expression (Lee et al., 2012; N. Liu & Stephanopoulos, 2021; Stephanopoulos, 2012). While useful, these methods are time-consuming and have not always been effective at improving performance, in part because these engineered strains are constructed sequentially in a low-throughput manner (Nielsen & Keasling, 2016). Furthermore, iterative introduction of single genetic changes does not explore the full space of possible combinatorial effects due to the inherent size and complexity of metabolic interaction networks, ultimately reaching a plateau (Hackett et al., 2016; Yang et al., 2018). This space of possibilities could be expanded by applying tools for target prediction such as *in silico* metabolic modelling, which lately is boosted by machine learning algorithms, for genotype-phenotype profiling (Lewis et al., 2012; Lu et al., 2019). Statistical modelling frameworks such as design of experiments have also been developed to facilitate probing the gene expression space and expedite the engineering of overproduction phenotypes (P. Xu et al., 2017; Yilmaz et al., 2022).

Random strain engineering methods augment prediction-based approaches by generating new hypotheses and exploring novel genetic space previously inaccessible. Following the principles of Darwinian evolution, random metabolic engineering methods rely on iterating rounds of genetic changes with screening or selection to identify strains with the desired fitness level, in this case productivity (Cao et al., 2020). These random genetic alterations could be introduced by methods such as adaptive lab evolution (ALE) (Sandberg et al., 2019), as well as chemical (Leavitt et al., 2017) and UV mutagenesis (Wang et al., 2020). Despite their success at improving strain performance, caveats of random genome evolution methods include sampling only a small fraction of the space of possible evolutionary trajectories, and the need for whole-genome sequencing to identify beneficial changes, combined with laborious genotype to phenotype mapping (Wang et al., 2020). This could be overcome by using other genome evolution methods that offer targeted and traceable approaches, such as the synthetic chromosome rearrangement and modification by loxP-mediated evolution (SCRaMbLE) (Dymond et al., 2011), RNAi Assisted Genome Evolution (RAGE) (HamediRad et al., 2018), and eukaryotic multiplex automated genome engineering (eMAGE) (Barbieri et al., 2017). As well, the clustered regularly interspaced short palindromic repeats (CRISPR)/CRISPR-associated (Cas) system has revolutionized targeted genome engineering and evolution. In addition to genome wide gene disruption libraries (Bao et al., 2015, 2018), gene activation and interference libraries have been created using catalytically dead Cas (dCas) protein variants that serve specific regulatory functions (Gilbert et al., 2014; Konermann et al., 2014). Another successful example is the strategy for CRISPR-mediated bidirectional titration of gene expression, which separately targeted subsets of the genome of *Saccharomyces cerevisiae* (Bowman et al., 2020). Genome evolution using CRISPR systems was also implemented through the creation of a trifunctional CRISPR system for the introduction of simultaneous alterations of differing nature *i.e.*, activation, interference, and deletion of multiple target genes (Lian et al., 2017). This trifunctional CRISPR system was then developed into a comprehensive multi-functional genome-wide CRISPR (MAGIC) system targeting every gene of the *S. cerevisiae* genome, which was used to engineer strains with enhanced tolerance to the toxic compound furfural (Lian et al., 2019). Although very efficient, the application of these methods for host metabolism optimization has two main drawbacks. First, single perturbation screens only capture a fraction of the best strategies possible, as metabolic flux rewiring is most effective when multiple adjustments are implemented simultaneously (Wang et al., 2020; J. Wu et al., 2015; M. Y. Wu et al., 2017). Second, the lack of a measurable phenotype in many cases necessitates the development of high-throughput methods given the large size of genome interrogation libraries.

Allosteric transcription factor (aTF)-based biosensors are useful tools for regulating promoter activity in response to specific small molecules (De Paepe et al., 2017; Fernandez-López et al., 2015), and have been adopted in metabolic engineering for detecting pathway intermediates and end products, metabolic flux monitoring, and dynamic regulation (Mahr & Frunzke, 2016). As well, aTF-based biosensors have proven effective for screening for overproduction phenotypes in strains modified by random genome engineering (Leavitt et al., 2017; Sandberg et al., 2019; Wang et al., 2020). This is in part due to the facile application of biosensors, where fluorescence or antibiotic resistance reporters could be used for rapid high-throughput screening or selection of large cell populations (Yuan & Ching, 2015). Furthermore, advances in protein engineering enabled modulating the ligand specificity and dynamic range of aTF biosensors (Nasr et al., 2023; Wan et al., 2019).

In this work, we highlight the utility of aTF-based biosensors as high-throughput screening and selection tools when combined with the MAGIC genome evolution tool (Lian et al., 2019) for optimizing shikimate pathway-derived *cis*,*cis*-muconic acid (ccM) production in *S. cerevisiae* (Pyne et al., 2018; Weber et al., 2012). Using this industrially relevant polymer precursor as a case study, we start by constructing a suite of functional biosensors for intermediates of the ccM pathway. We then develop a population enrichment method leveraging the zeocin resistance marker *bleoR* or GFP as biosensor reporters to be used for library interrogation. Building on this, we employ the biosensor-directed MAGIC (senMAGIC) framework to identify genetic determinants that improve strain productivity in two strain backgrounds, one probing ccM productivity directly, and the other indirectly through PCA (the first committed pathway intermediate) as a proxy for shikimate pathway flux. Finally, we combine the top hits from this screen to build a multiplexed gRNA library targeting multiple genes simultaneously and identify both intuitive and novel changes that increase ccM productivity, confirming the top hits by implementing the changes directly into CRISPR-free production strains.

## RESULTS

### Biosensor Development and Validation

Prokaryotic allosteric transcription factor (aTF)-based biosensors have been transplanted into eukaryotic organisms such as the yeasts *S. cerevisiae* and *Yarrowia lipolytica* (Lv et al., 2020; Skjoedt et al., 2016), as well as to mammalian cell-lines (Xie & Fussenegger, 2018). To aid in the metabolic engineering of the ccM production pathway, we identified candidate aTFs targeting pathway intermediates, namely protocatechuate (PCA), catechol, ccM, and the final pathway molecule adipic acid (Fig. 1a). We characterized the response of each biosensor by measuring fluorescence of a GFP reporter under aTF regulatory control in the presence or absence of its inducer. Successful induction of expression was observed from the PcaV-regulated promoter by PCA, the Caq101-regulated promoter by catechol, the BenM^WT^- regulated promoter by ccM, and the BenM^Tisno^-regulated promoter by adipic acid (Fig. 1b, c, Fig. S1d) (Skjoedt et al., 2016; Snoek et al., 2020).

**Fig. 1.**
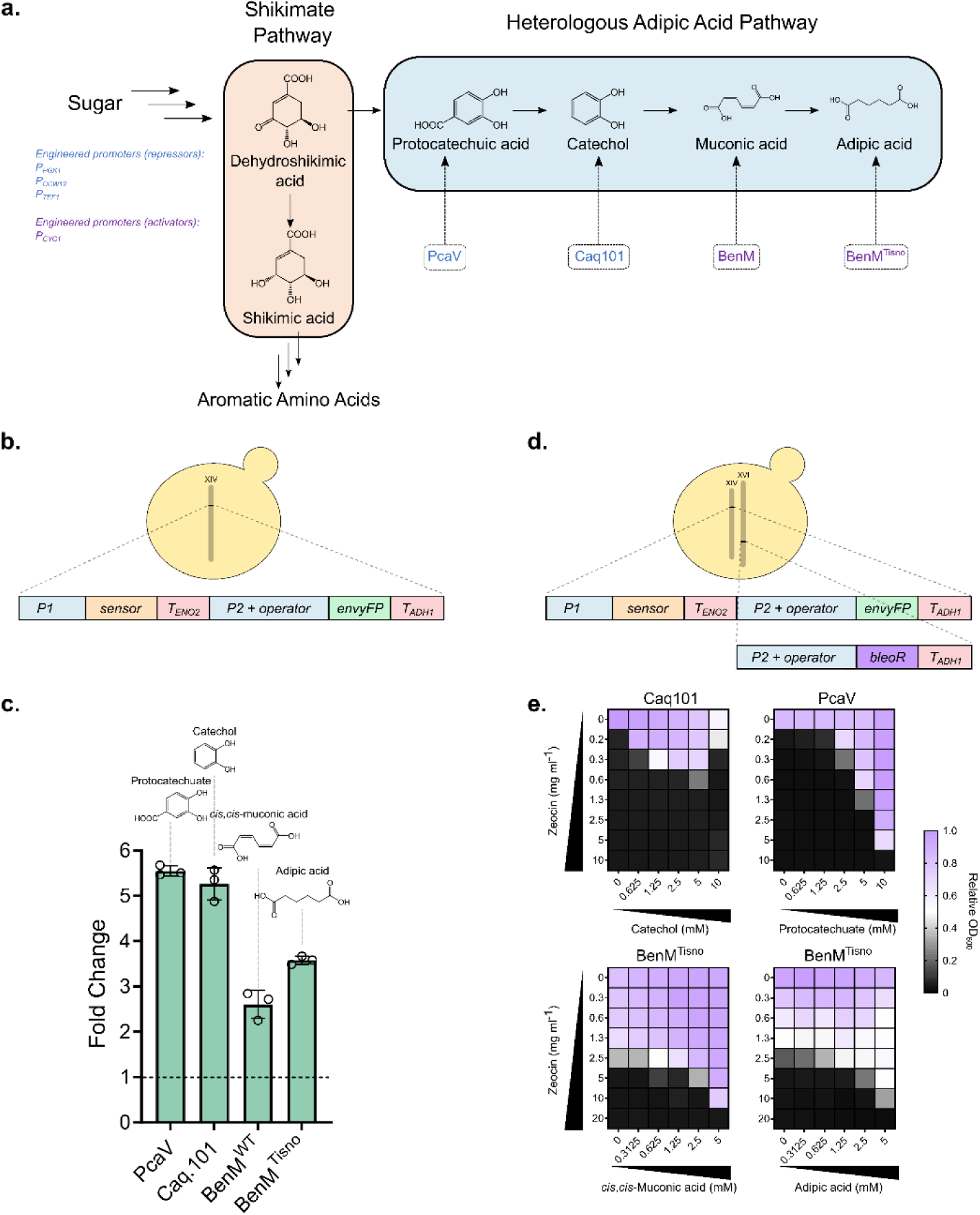
ccM pathway and biosensor design. **a** The shikimate pathway-derived ccM pathway in *S. cerevisiae* and repressor (blue) and activator (purple) biosensors. Promoter(s) engineered in this study for repressors (blue) and activators (purple) are listed. **b** Biosensor cassette integrated into *S. cerevisiae*. **c** Fluorescence response of biosensor candidates to their cognate inducers at a concentration of 5 mM. Datapoints are mean ± SD, *n* = 3. **d** Biosensor and zeocin resistance cassettes integrated into *S. cerevisiae*. **e** Double gradient of different concentrations of zeocin and the inducer for Caq101 with catechol, PcaV with protocatechuate, BenM^Tisno^ both with ccM and adipic acid. Datapoints are mean, *n* = 2.

To optimize the biosensor promoters, we tested additional promoter architectures for each of the repressors, PcaV and Caq101, using the promoters P*_CCW12_*, *P_PGK1_*, and *P_TEF1_*. In the presence of catechol, the Caq101-regulated *P_CCW12_* promoter variant had a superior induction of up to ∼9 fold compared to the Caq101-regulated *P_PGK1_*and *P_TEF1_* variants, which had a maximum fold induction of ∼2 and ∼7 fold, respectively (Fig. S1a). For PCA, we found all three PcaV-regulated promoters to have an equivalent fold-change response, albeit being leaky (Fig. S1b, S2). We reduced this leakiness and improved dynamic range by introducing a second operator copy immediately downstream of the first in the PcaV-regulated *P_CCW12_* variant, increasing fold-change by up to 15-fold (Fig. S1c, S2).

For ccM detection, the BenM-regulated *P_CYC1_* promoter achieved ∼2.5 fold induction in the presence of ccM (Fig. S1d). The BenM^Tisno^-regulated *P_CYC1_* promoter, which is leakier than the BenM^WT^-regulated variant, had higher maximal fold inductions of ∼7 and ∼9 fold by ccM and adipic acid, respectively (Fig. S1d, S2). Crosstalk analyses of PcaV, Caq101, BenM^WT^, and BenM^Tisno^ showed the specificity of PcaV, BenM^WT^, and Caq101 to their cognate inducers PCA, ccM, and catechol, respectively. However, the BenM^Tisno^ variant had notable crosstalk, responding to ccM, adipic acid, and catechol (Fig. S1e). Based on these results, we selected the PcaV and BenM^Tisno^ biosensors for further development, as they provided the most specific or strongest responses to PCA and ccM, the key intermediates of interest in our pathway.

### Expanding biosensors into population enrichment tools and metabolite reporters

To develop a selectable biosensor system, we introduced the zeocin resistance marker *bleoR* as a reporter under the control of the aTF-regulated *P_CCW12_* or *P_CYC1_* promoters into the respective biosensor strains, mirroring the GFP reporter design (Fig. 1d). Zeocin, a phleomycin-family glycopeptide antibiotic, induces double-stranded DNA breaks in a dose-dependent manner, with higher concentrations selecting for increased *bleoR* expression, as BleoR inactivates zeocin stoichiometrically (Chankova et al., 2007; Hauber et al., 1985; Yuan & Ching, 2015; Zhao et al., 2018, 2019). Due to variable basal expression, we anticipated that biosensor leakiness would influence zeocin resistance, with promoters regulated by PcaV and Caq101 being the least leaky, and by BenM^Tisno^ being the most (Fig. S2). As expected, titrating both inducer and zeocin concentrations revealed a graded cellular growth response: PcaV and Caq101 strains were sensitive to zeocin in the absence of inducers but gained resistance when PCA or catechol were supplied, with growth inhibition observed in high catechol concentrations due to its toxicity (Liao et al., 2024) (Fig. 1e). In contrast, the uninduced BenM^Tisno^ strain exhibited growth up to 2.5 mg mL^-1^ zeocin, increasing to 10 mg mL^-1^ in the presence of ccM or adipic acid (Fig. 1e).

Given the role of PCA as the first committed intermediate in the ccM pathway (Fig. 1a), we next evaluated whether the PcaV biosensor could function as a reporter of shikimate pathway flux towards heterologous production of PCA and ccM. To generate PCA endogenously, we introduced the dehydroshikimate dehydratase *AROZ* from the fungus *Podospora anserina* into the PcaV sensor strain, enabling conversion of dehydroshikimate to PCA (Fig. 2a). This modification resulted in ∼8-fold GFP induction compared to a strain lacking *AROZ*, with a similar trend observed in PCA titres (Fig. 2a, c). Further testing showed that supplementing phenylalanine or tyrosine—feedback inhibitors of native DAHP synthases Aro3p and Aro4p (Paravicini et al., 1989)—reduced both fluorescence and PCA levels (Fig. 2c), validating the system as a flux-responsive reporter for PCA production via the shikimate pathway. In parallel, introducing the BenM^Tisno^ sensor into a ccM-producing strain increased GFP fluorescence by ∼2.5-fold relative to a control strain lacking the pathway, reflecting the ccM productivity of both strains (Fig. 2b, d) Together, these results demonstrate that the PcaV and BenM^Tisno^ biosensors not only enable zeocin-based selection but also serve as functional reporters of flux through the shikimate pathway towards PCA and ccM. Both PCA and ccM strains described here serve as parents for M1 and M3, respectively, described in the following section (Fig. 2a, b).

**Fig. 2.**
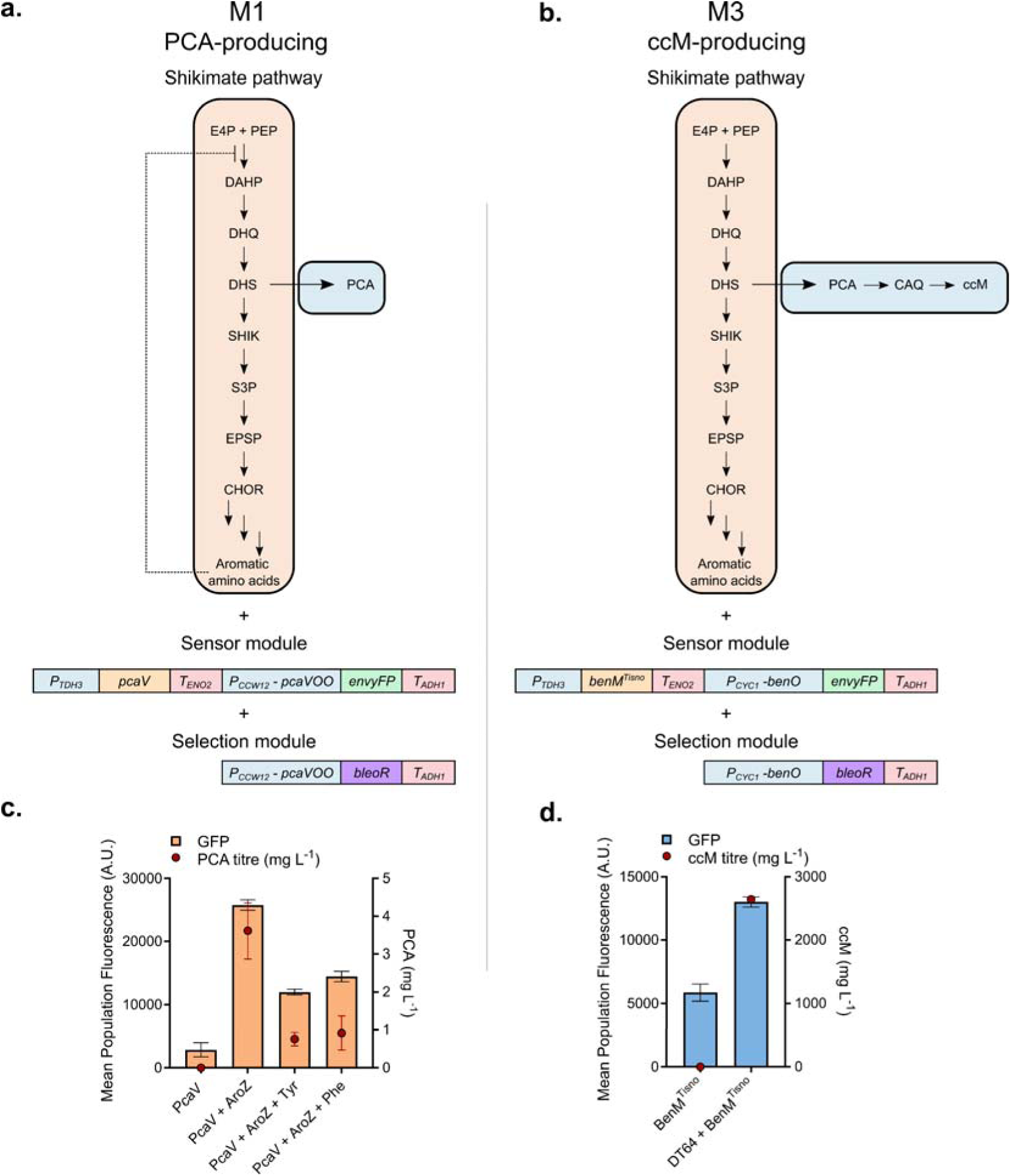
Using biosensors for measuring endogenously produced PCA and ccM. **a** PCA production strain carrying AroZ (dehydroshikimate dehydratase) in combination with the PcaV sensor and zeocin resistance cassettes. **b** ccM production strain encoding the full ccM pathway and an abolished feedback inhibition in the shikimate pathway, in combination with the BenM^Tisno^ sensor and zeocin resistance cassettes. **c** Biosensor response and PCA titres of strains carrying AroZ and the PcaV biosensor grown in minimal YNB medium with or without tyrosine or phenylalanine supplementation. **d** Biosensor response and ccM titres (conducted as separate experiments) of strains carrying the BenM^Tisno^ sensor in a wildtype background and in the ccM production strain DT64. Datapoints are mean ± SD, *n* = 3. E4P Erythrose-4-phosphate, PEP Phosphoenolpyruvate, DAHP 3-Deoxy-D-arabino-heptulosonate-7-phosphate, DHQ 3-Dehydroquinate, DHS 3-Dehydroshikimate, SHIK Shikimate, S3P Shikimate-3-phosphate, EPSP 5-Enolpyruvylshikimate-3-phosphate, CHOR Chorismate, PCA Protocatechuic acid, CAQ Catechol, ccM *cis*,*cis*-muconic acid.

### senMAGIC Screen for Improving ccM pathway productivity

To test the senMAGIC framework for biosensor-assisted screening of genome-wide gRNA libraries, we constructed two fit-for-purpose strains that interrogate distinct aspects of the ccM pathway. Strain M1 was built to identify changes that improve shikimate pathway flux and is based on the strain carrying both AroZ and the PcaV biosensor with dual BleoR and GFP reporters (Fig. 1e-2a). Strain M3 was built to identify changes that increase ccM titres. This strain comprises the BenM^Tisno^ biosensor with dual BleoR and GFP reporters (Fig. 1e, 2b) in the ccM-producing background DT64, which carries genomically integrated copies of the ccM pathway enzymes and harbours a degron-tagged Aro1p combined with abolished shikimate pathway feedback inhibition as described in our previous work (Pyne et al., 2018). Strains M1 and M3 additionally carry a *URA3* deletion to enable auxotrophic selection of gRNA library plasmids and a genomically-integrated trifunctional CRISPR plasmid to facilitate gene activation, interference, or deletion (Lian et al., 2019).

Next, we aimed to study genetic determinants that enhance flux through these pathways using a CRISPR-AID genome-wide library screen (Lian et al., 2019). M1 and M3 strains transformed with the activation, interference, and deletion gRNA libraries were grown in zeocin to enrich genetic perturbations that resulted in increased PCA or ccM production, followed by gRNA sequence analysis by next-generation sequencing (Table S6). Among the most enriched gRNAs, some expected candidates were found for M1, such as the chorismate synthase *ARO2* interference gRNA, which likely attenuates flux down the shikimate pathway towards amino acid production to be diverted towards PCA production. In contrast, the M3 gRNA sequence analysis did not return any hits with effects that could be rationalized. For this reason, we selected the top 100 most enriched gRNA hits for M1 and expanded the selection to include the top 200 hits for M3 to be used for subsequent experiments. Additionally, since zeocin is a DNA-damaging agent (Chankova et al., 2007), we omitted any gRNAs with obvious DNA-repair function to avoid false-positive hits (Table S7 and S8).

### Combinatorial CRISPR-AID Library Design and QC

Single-target approaches in metabolic engineering face a bottleneck in efficacy improvements due to the complexity of genetic and metabolic networks, motivating the use of multiplexed perturbation strategies (Wang et al., 2020; J. Wu et al., 2015; M. Y. Wu et al., 2017). We therefore constructed a combinatorial CRISPR-AID library using the identified hits as a template, with the aim of uncovering additive or synergistic changes that increase PCA and ccM titres.

For practical screening purposes, we limited the combinatorial space to three gRNAs per plasmid and used the tRNA_GLY_ splicing units previously developed for multiplexed gRNA expression as a single transcript (Zhang et al., 2019). We first verified the functionality of the design by comparing the activation, interference and deletion with gRNAs expressed alone or in a triple-gRNA array spliced by two tRNA_GLY_ units (Fig. S3). Strains carrying the triple-gRNA array simultaneously exhibited the three types of perturbations, albeit at a lower level compared to the single-gRNA design (Fig. S3). We next designed a cloning method to randomly combine three gRNAs from the top hits previously selected. Hits from the single-gRNA library were PCR-amplified with specific primers and used to construct gRNA array plasmids in *Escherichia coli* using Golden Gate assembly (Fig. S4 and supplementary method 1). Combining 100 gRNAs for M1 and 200 gRNAs for M3 in arrays of 3 gRNAs, we aimed for a minimum number of clones of 100^3^ = 10^6^ and 200^3^ = 8.10^6^ for M1 and M3 libraries, respectively (see supplementary method 1 for detailed cloning data).

For quality control of the library, plasmids from 96 isolates per library were PCR-amplified with barcoded primers, pooled, and sequenced (see supplementary method 2 for details). Analysis of the sequencing data showed that the cloned library had gRNA distributions that drifted from the original ordered library, with activation gRNAs being over-represented in M1, and underrepresented in M3 (Fig. S5a). The numbers of gRNAs per array ranged from one to four, with more than half arrayed plasmids containing three gRNAs in both M1 and M3 libraries (Fig. S5b). Over-represented gRNAs were rare; however, the few gRNAs that were over-represented were found 20-60 times in the population (Fig. S5c).

### Combinatorial Library Screening and Demultiplexing

To identify multiplexed determinants that improve PCA and ccM production, we transformed the combinatorial gRNA libraries into strains M1 and M3 and subjected the transformants to zeocin selection, followed by FACS to select the top GFP signaling cells (Fig. 3a). We then analyzed the productivity of 95 colonies per sorted library (Fig. 3b) and sequenced the gRNAs from the best producers (Table 1).

**Fig. 3.**
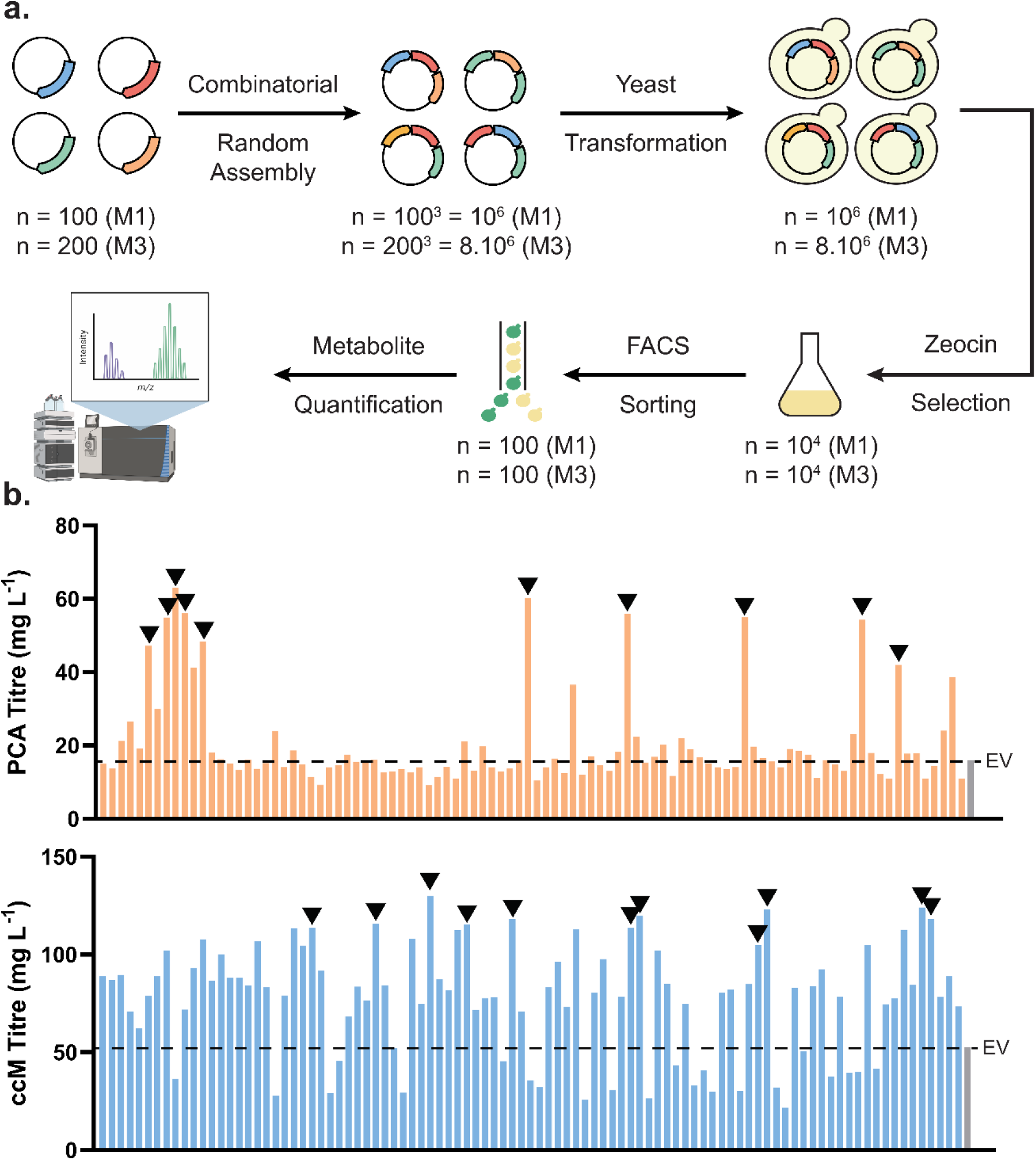
CRISPR-AID combinatorial screen. **a** General workflow of the combinatorial screen. **b,c** PCA (b) and ccM (c) titres of the 95 best candidates obtained after zeocin selection and screened with FACS and GFP fluorescence signal. Grey bar denotes the empty vector (EV) strain. Black arrows highlight the best hits selected for sequencing. Part of figure 3a made with Biorender.com. PCA Protocatechuic acid, ccM *cis*,*cis*-muconic acid.

**Table 1:** Best hits performances and gRNA targets and activities (a=activation, i=interference) from M1 and M3 combinatorial screen.

| **Colony name** | **PCA titre (mg.L<sup>-1</sup>)** | **PCA yield (mg.L<sup>-1</sup>.OD<sup>-1</sup>)** | **gRNA 1** | **gRNA 2** | **gRNA 3** |
|---|---|---|---|---|---|
| M1_EV | 22.07 | 31.26 | /* | / | / |
| M1_A09 | 62.97 | 76.54 | SOE1i | RVS167i | / |
| M1_D12 | 60.14 | 90.50 | DUG3i | PDR3a | RNH202a |
| M1_A10 | 56.09 | 99.32 | CCS1a | RIM4i | PDR3a |
| M1_E11 | 55.99 | 106.94 | PAU19a | RUP1i | DML1_a |
| M1_F12 | 54.99 | 86.88 | ATG32a | SNR79a | OYE3a |
| M1_A08 | 54.90 | 82.62 | ND | ND | ND |
| M1_H01 | 54.30 | 87.55 | ND | ND | ND |
| M1_A12 | 48.29 | 74.17 | LCB5i | TGL3a | TOS6a |
| M1_A06 | 47.15 | 66.45 | MIC27i | KRE28a | NIS1a |
| M1_H05 | 41.99 | 93.94 | ATG32a | TGL3a | BCK1d |
| **Colony name** | **ccM titre (mg.L<sup>-1</sup>)** | **ccM yield (mg.L<sup>-1</sup>.OD<sup>-1</sup>)** | **gRNA 1** | **gRNA 2** | **gRNA 3** |
| M3_EV | 52.62 | 7.66 | / | / | / |
| M3_D01 | 130.11 | 14.60 | TPI1i | TPI1i | SNR60i |
| M3_H07 | 124.12 | 13.60 | RIX7i | TPI1i | TUB3i |
| M3_G02 | 122.90 | 13.88 | IDP1i | / | / |
| M3_E12 | 119.65 | 11.79 | IDP1i | / | / |
| M3_H08 | 118.27 | 11.10 | IDP1i | / | / |
| M3_D10 | 118.24 | 11.83 | IDP1i | / | / |
| M3_C07 | 115.72 | 14.07 | ATX1i | LHS1a | TPI1i |
| M3_D05 | 115.31 | 15.77 | TIM54i | TPI1i | / |
| M3_E11 | 113.81 | 14.23 | ATX1i | TPI1i | TPI1i |
| M3_B12 | 113.70 | 11.18 | IDP1i | / | / |
| M3_G01 | 104.83 | 22.38 | TPI1i | / | / |

\* No guide present at this position in the gRNA array.
\*\* ND: Not Determined. Guides could not be identified from sequencing data for those hits.

In strain M1, a diverse set of gRNAs that resulted in increased PCA titres emerged from the screen, and a few repeated hits were observed. Indeed, only one gRNA activating the *ATG32* gene involved in mitophagy was found twice across the screen (Table 1). The direct mechanism of *ATG32* activation and the other enriched gRNAs for M1 lack clear links to the shikimate pathway, suggesting that improvement may stem from broader metabolic or regulatory changes influencing PCA titres. In contrast, the M3 screen yielded more consistent hits that improved ccM titres, with strong enrichment of interference gRNAs targeting *TPI1* and *IDP1*. *TPI1* encodes a triose phosphate isomerase, which catalyzes the interconversion of dihydroxyacetone phosphate (DHAP) into glyceraldehyde phosphate (GAP). As DHAP is a glycolysis intermediate that undergoes a series of reactions to be converted to the shikimate pathway precursor phosphoenolpyruvate (PEP) (Fig. 2a, b), alterations in its cellular abundance could influence carbon flux towards ccM. *IDP1* encodes a NADP-dependent isocitrate dehydrogenase not involved in the tricarboxylic acid cycle despite its mitochondrial location. Although the effect of this perturbation on ccM productivity is less clear, it is strongly selected from the screen where it appears in 5 of the top 11 hits and enhances ccM titres by 2.6-fold to a final concentration of 122.9 mg L^-1^, whereas the *TPI1* interference resulted in a 2.2-fold titre increase (Table 1).

Intriguingly, we could not analyze the sequencing data for some of the best hits of M1 (Table 1). Triple-gRNA plasmid extraction from a yeast clone followed by transforming into *E. coli* and sequencing multiple bacterial colonies yielded different gRNA sequences, strongly suggesting that more than one plasmid was present within the same yeast clone (Fig. S6). This is possibly due to the high plasmid concentration used for the transformation of large combinatorial libraries, leading to multiple plasmids being present in a yeast cell (Wäneskog et al., 2025). This artefact was not observed for the M3 best hits, possibly because longer growth during zeocin exposure allowed better plasmid segregation.

We next investigated the contribution of individual gRNAs from the top three PCA-producing colonies of strain M1, and the top two ccM-producing colonies of strain M3 identified in the combinatorial screen (Table 1). For M1 gRNAs, we reconstructed plasmids that test the full cassette identified in the screen, as well as individual gRNAs and their different combinations (Fig. 4a). For M3 gRNAs, we chose colonies B12 and G01 as they were the most enriched in our screen and resulted in high improvements in ccM productivity (Table 1). Since these M3 plasmids comprised single gRNAs targeting *IDP1* and *TPI1*, we combined them to analyze their combinatorial effect (Fig. 4b).

**Fig. 4:**
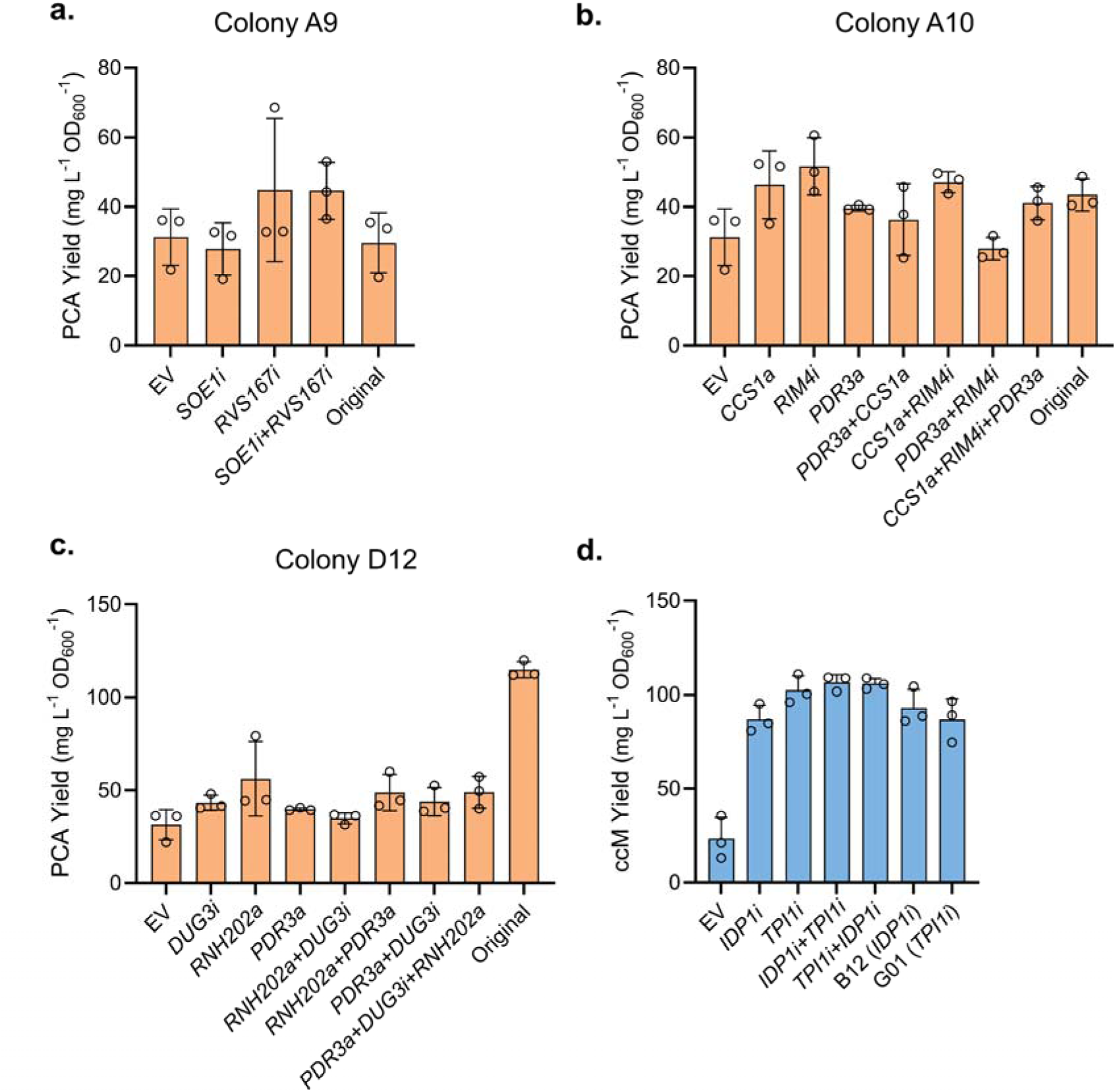
Combinatorial breakdown of M1 & M3. **a-d** Top three hits from the PCA screen (a-c) and ccM screen (d) were sequenced and deconvoluted. The guide array plasmids were rebuilt with corresponding single, double, and/or triple guides and grown alongside the original colony obtained from the screen for PCA or ccM production, respectively. Datapoints are mean ± SD, *n*= 3 biological replicates. PCA Protocatechuic acid, ccM *cis*,*cis*-muconic acid.

On building the different M1 combinations, we observed that colony A9 did not reproduce the original increase in PCA yields observed in the screen (Fig. 4a). For colony A10, *CCS1* activation and *RIM4* interference each increased PCA yields by ∼50% and 65%, respectively, with no additive effect observed in combination (Fig. 4b). *PDR3* activation resulted in a ∼30% increase in PCA yields, with a similar trend of no observed additivity when combined with other gRNAs, and a decrease in productivity by ∼10% when combined with a *RIM4* interference gRNA (Fig. 4b). No appreciable increase was seen for colony D12 with any gRNA or combination. However, both the original and cured D12 strain (retransformed with an empty vector) showed elevated PCA production of 93% and 72%, respectively, suggesting a strain-specific background effect independent of the gRNAs (Fig. 4c, S7). For M3 combinations, interference gRNAs targeting *IDP1* and *TPI1* individually increased specific ccM production by 274% and 340% respectively, compared to the empty vector control (Fig. 4d). Combining *IDP1* and *TPI1* interference did not yield an additional benefit over *TPI1* interference alone. Furthermore, reversing the position of the *IDP1* and *TPI1* gRNAs within the multiplexed cassette did not affect ccM production (Fig. 4d).

### senMAGIC hit validation by genomic integration

Genome-wide CRISPR systems and plasmid-based screens in yeast are inherently prone to false positives and negatives (Bock et al., 2022). This challenge is compounded by the multiple regulatory layers applied in our senMAGIC workflow. To confirm the true effects of the perturbations identified in our screen, we engineered a subset of the genetic modifications directly into CRISPR-free strains that produce PCA and ccM and analyzed them in microtiter plate fermentations (Fig. 2a, b). For activation and interference, we introduced a strong (*P_TDH3_*) or weak (*P_CYC1_*) *S. cerevisiae* promoter, respectively, upstream of the genes of interest (Apel et al., 2017) (Fig.5a). *P_CYC1_*was truncated to remove its upstream activating sequences for a further reduction in expression (Skjoedt et al., 2016).

For the single gRNA senMAGIC screen, inserting the truncated *P_CYC1_*variant immediately upstream of *ARO2* in a PCA-producing strain resulted in a ∼100% increase in PCA yields (Fig. 5b). Introducing this modification in a ccM strain resulted in a modest 12% increase in ccM yields, possibly as this strain harbours a degron-tagged ARO1p that limits flux down the main branch of the shikimate pathway and increases productivity (Fig. 5c) (Pyne et al., 2018)

**Fig. 5:**
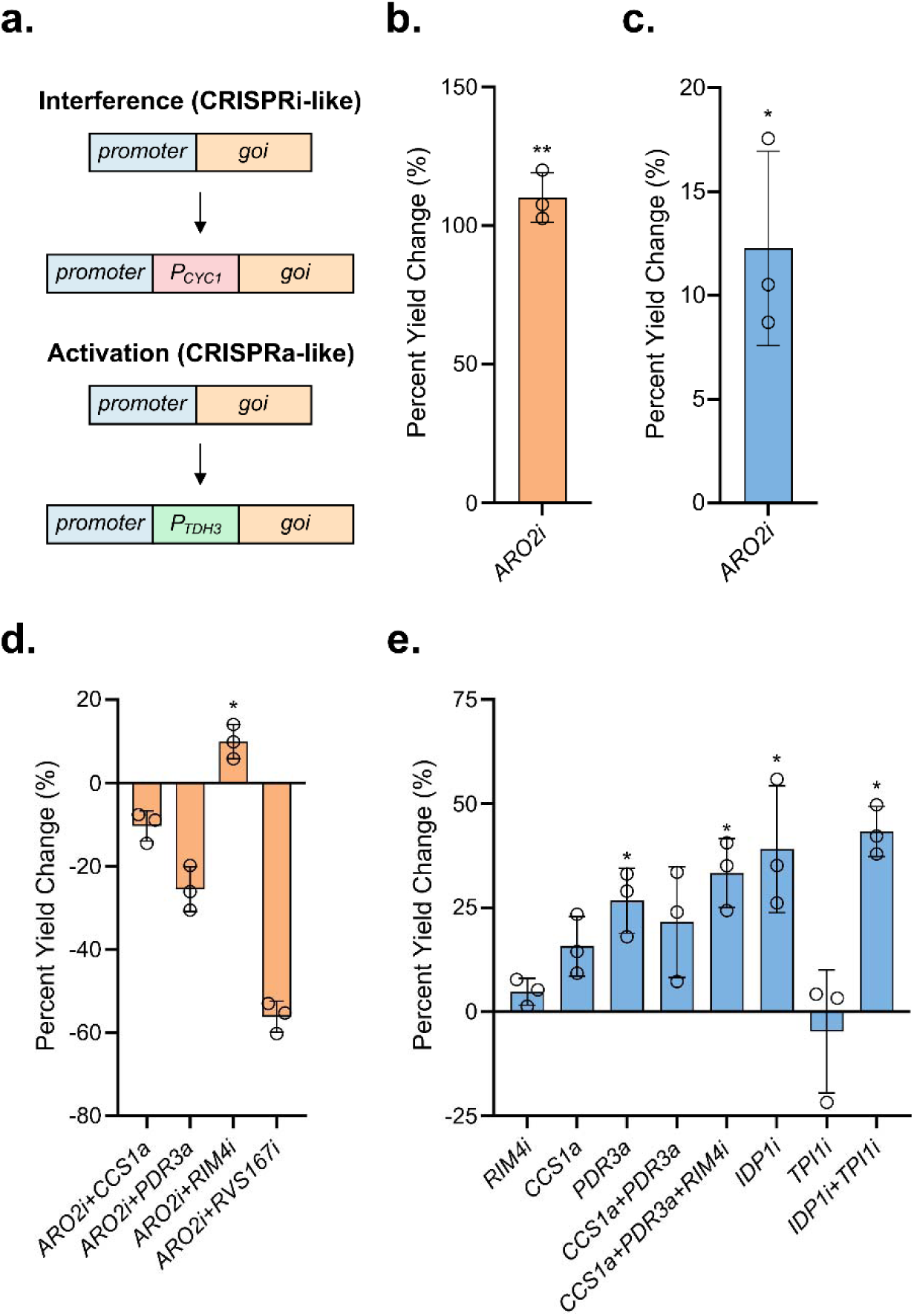
Validation of senMAGIC hits by genomic integration into CRISPR-free strains. **a** Schematic for strain construction methodology for interference and activation. **b.** PCA percent yield change after introducing a weak promoter for *ARO2* in a PCA-producing strain compared to a strain carrying the wild-type promoter. **c.** ccM percent yield change after introducing a weak promoter for *ARO2* in a ccM-producing strain compared to a strain carrying the wild-type promoter. **d.** PCA percent yield change after promoter replacements for *CCS1*, *PDR3*, *RIM4*, and *RVS167* in a PCA-producing strain compared to a strain carrying the wild-type variants of these promoters. **e.** ccM percent yield change after promoter replacements for *CCS1*, *PDR3*, *RIM4*, *RVS167*, *IDP1*, and *TPI1* alone or in combination. All promoter replacements were conducted in PCA- or ccM-producing strains and compared so the same strains carrying wild-type variants of these promoters to calculate percent yield change. Datapoints are mean ± SD, *n* = 3. Statistical significance was determined by a two-tailed Student’s t-test by comparing strain titres to the respective parent producer strain; **P < 0.01; *P < 0.05. PCA Protocatechuic acid, ccM *cis*,*cis*-muconic acid.

In a PCA-producing strain carrying the *ARO2* promoter modification, adding the strong *P_TDH3_* promoter upstream of *CCS1* and *PDR3* or *P_CYC1_* upstream of *RVS167* and *RIM4* resulted in a drop in strain productivity in all cases with the exception of *RIM4*, which further improved yields by 15% (Fig. 5d). In the ccM-producing strain, inserting *P_TDH3_* promoter upstream of *CCS1* and *PDR3* resulted in ccM yield increases of 15% and 26%, respectively, and 21% when both modifications were combined in the same strain (Fig. 5e). Introducing *P_CYC1_* upstream of *RIM4* resulted in a minor 5% increase, but when combined with the *CCS1* and *PDR3* modifications resulted in a ∼33% ccM yield increase (Fig. 5e). Finally, inserting *P_CYC1_* upstream of *IDP1* resulted in a ∼39% yield increase, but introducing the same modification upstream of *TPI1* resulted in a drop in productivity, suggesting that precisely tuning *TPI1* expression might be needed. Combining both *IDP1* and *TPI1* modifications resulted in a ∼40% increase in yields (Fig. 5e).

A preliminary assessment of different carbon sources (glucose, sucrose, slow-release) and fermentation duration showed that fermentations of the PCA-producing strain carrying the *ARO2* promoter modification in a slow-release carbon source for 72 hours markedly improved PCA yields (Fig. s8). No appreciable increase was observed in the ccM strain carrying the same modification. These results suggest that further productivity improvements in strains carrying senMAGIC-identified perturbations relative to the parent strain could be achieved through optimization of media composition and fermentation conditions.

## Discussion

Microbial biocatalysis provides the basis for sustainable production of value-added chemicals to reduce fossil fuel dependency, including PCA and ccM, precursors to valuable platform chemicals. In the present study, we combined a biosensor-directed population selection method with a genome-wide CRISPR screen to identify genetic determinants that increase PCA and ccM titres in yeast. Using this multi-functional CRISPR system that targeted all ∼6000 genes of the *S. cerevisiae* genome for activation, interference, and deletion, we identified several intuitive and non-intuitive targets that improved strain productivity. We then used the top single hits to build combinatorial libraries that uncover possible additive and synergistic effects. As a proof of concept for this approach, implementing these modifications resulted in maximum titre improvements of ∼110% and ∼40% for PCA and ccM, respectively, which could be further boosted by prolonging fermentation duration or altering carbon sources.

Orthogonal control of gene expression is one of the hallmarks of synthetic biology (Medema et al., 2011). aTF-based biosensors are excellent tools for controlling gene expression owing to their diversity of chemical inducer signals, simplicity of heterologous expression as small soluble proteins, and their portability and functionality in eukaryotic systems (Fang et al., 2016; Ikushima et al., 2015; Lv et al., 2020; Matilla et al., 2022; Snoek et al., 2020). Here, we took advantage of this inherent diversity to engineer and assess the responsiveness of four biosensor candidates targeting each metabolite in the ccM/adipic acid pathway in *S. cerevisiae*. To select for high-producing strains in a mixed population, we extended the biosensor reporter beyond fluorescence output to include the zeocin resistance marker *bleoR* (Tominaga et al., 2021). Zeocin is a dose-dependent antibiotic, which enables selection for expression strength of *bleoR* (Yuan & Ching, 2015; Zhao et al., 2019). This feature allowed us to account for biosensor promoter leakiness, which we observed in zeocin inhibitory concentration differences between the tight repressors Caq101 and PcaV, and the leaky activator BenM^Tisno^. Furthermore, zeocin inhibitory concentrations increased with all three biosensors upon exogenously supplementing their cognate aTF inducers. Nevertheless, a notable caveat of using zeocin is its ability to introduce DNA lesions such as double-stranded breaks, with a preference for the five-base sequence 5′-TGTNC-3′ (Zheng et al., 2022). In earlier iterations of this work, zeocin-induced DNA lesions combined with the *S. cerevisiae* propensity for homology-directed repair (Storici et al., 2003) resulted in the inactivation of the functional copy of *PAD1* by introducing a mutation that matched the native CEN.PK *PAD1* allele, abolishing ccM titres (Pyne et al., 2018). This could be corrected by ensuring minimal homology between sequences in engineered strains to eliminate inadvertent DNA donor templates that could be used for zeocin-induced homologous recombination, a consideration of broad relevance when using zeocin in metabolically engineered hosts (Zheng et al., 2022). Additionally, following this workflow, some strains exhibited high titres that persisted after gRNA plasmid curing, suggesting the possibility of unknown effects resulting from exposure to zeocin. This necessitated confirmation of the genetic changes identified in our workflow by genomic integration in strains not exposed to zeocin.

aTF-based biosensors can be engineered to monitor metabolic flux in a given engineered pathway (Li et al., 2017; P. Xu, 2018), and coupling biosensors with enzymes further expands the repertoire of detectable molecules (Boada et al., 2020; Bowman et al., 2021). We applied these principles by incorporating the BenM^Tisno^ ccM biosensor directly in a ccM-producer strain as well as engineering a shikimate pathway flux biosensor, for which we lacked a dedicated aTF in our collection, by coexpressing the dehydroshikimate dehydratase AroZ alongside the PcaV biosensor. This is a useful and generalizable strategy for metabolic engineering in the absence of dedicated biosensors for the molecule(s) of interest. Additionally, promoter strength driving AroZ expression serves as a tunable parameter to optimize sensor dynamic range, as strong promoters risk saturating the biosensor at low flux rates, limiting its utility.

CRISPR-mediated genome-interrogation methods can be used for engineering complex phenotypes either by targeting the entire genome or subsets with specific metabolic functions *e.g.*, genes involved in carbohydrate and lipid metabolism or transporter-encoding genes (Bowman et al., 2020; Lian et al., 2019; Wang et al., 2021). These approaches have the power of identifying genetic determinants that could otherwise be missed by classical approaches.

Here, we employed the MAGIC genome-wide multifunctional CRISPR tool in the biosensor production strains built to interrogate distinct aspects of the pathway, namely shikimate pathway flux and ccM pathway titres, thereby creating the senMAGIC platform. Single-perturbation library plasmids were transformed into strains, subjected to zeocin enrichment, and analyzed by next-generation sequencing to identify the most-enriched gRNAs. The top hits from each strain were then isolated and cloned into multi-gRNA arrays using Golden Gate, which yielded arrays of variable length containing one to four gRNAs that affected combinatorial library coverage.

Using this approach, we identified several activation and interference genetic targets that we then validated by inserting strong or weak promoters immediately upstream of the genes of interest in CRISPR-free strains, leaving their native promoters intact to avoid disrupting potential bidirectional promoter activity (Apel et al., 2017; Z. Xu et al., 2009). Among the hits from the single-gRNA library screen in the PCA-producing strain M1, *ARO2* interference via integration of a weak promoter resulted in a ∼100% increase in PCA yields in a CRISPR-free strain. This is consistent with the effect of expressing truncated or degron-tagged Aro1p variants, which reduce flux through the shikimate pathway and reroute it toward the heterologous branch (Leavitt et al., 2017; Pyne et al., 2018). A more modest ∼10% increase in ccM yields reflects a limited benefit of *ARO2* interference in this strain that harbours a degron-tagged Aro1p (Pyne et al., 2018).

The combinatorial gRNA library screen in both M1 and M3 yielded several candidate perturbations, which we validated by demultiplexing followed by promoter replacement. Introducing a strong promoter to drive *CCS1* and *PDR3* expression improved ccM yields by ∼15% and ∼26%, respectively. CCS1p is a chaperone that is required for Cu^2+^-loading of the superoxide dismutase SOD1p, and its overexpression was shown to relieve oxidative stress and extend chronological lifespan when simultaneously overexpressed with *SOD1* (Harris et al., 2005). Notably, a strain carrying a *CCS1* deletion was identified in a chemogenomic screen to have reduced adipic acid tolerance (Fletcher et al., 2021), and both *CCS1* and *SOD1* appeared as deletion hits in a recent genome-wide CRISPR screen, collectively underscoring the importance of investigating the role Ccs1p and Sod1p play in the ccM pathway (Cachera et al., 2024). *PDR3* is a transcription factor that regulates multiple efflux pumps, including PDR12, whose deletion abolished and overexpression improved dicarboxylic acid tolerance, respectively (Fletcher et al., 2021; Pereira et al., 2019), suggesting a potential benefit of interrogating the *PDR3* regulon for ccM pathway improvements. Similarly, overexpression of efflux pumps in the non-model yeast *Kluyveromyces marxianus* enhanced dicarboxylic acid tolerance, underscoring the broad applicability of this strategy (Thornbury et al., 2025). Additionally, replacing the endogenous promoters of *RIM4* and *IDP1* with a weak *P_CYC1_* resulted in ∼5% and ∼39% increases in ccM yields, respectively. Rim4p is an RNA-binding protein involved in regulating sporulation, which although unrelated to the ccM pathway, is not the first instance cell division regulators appear in genome-wide engineering efforts, as noted previously with *CDC15* mutations and *FOB1* overexpression (Cachera et al., 2024; Wang et al., 2020). Idp1p is a mitochondrial isocitrate dehydrogenase whose role in improving ccM yields remains unclear, though studies show it to be slightly upregulated upon catechol exposure (Liao et al., 2024), and that its deletion increases cell chronological lifespan (McCormick et al., 2015). Interference with *TPI1*, a triose phosphate isomerase that acts upstream of the shikimate pathway, may affect concentrations of the E4P and PEP precursors (Islam et al., 2017) and results in enhanced oxidative stress tolerance (Ralser et al., 2007). However, ccM yields improved only from a plasmid-based *TPI1* interference but not upon promoter replacement, suggesting that further *TPI1* expression tuning may be required for yield improvement. The largest yield improvements were observed in combinatorial perturbations: activation of *CCS1* and *PDR3* combined with *RIM4* interference resulted in a ∼33% yield increase, while *IDP1* interference alone or combined with *TPI1* resulted in a ∼40% increase in yields. Together with targets identified in parallel studies, such as *ZWF1* and methionine biosynthetic gene deletions, these perturbations contribute to a growing repertoire of validated engineering targets for ccM pathway optimization (Cachera et al., 2024; Curran et al., 2013).

We identified three main bottlenecks for senMAGIC that should be carefully addressed when using this framework. First, the choice of biosensor is critical, with specificity, leakiness, and dynamic range being key determinants of screen performance. Second, ensuring the proper coverage of arrayed genome-wide gRNA libraries in the cloning and plasmid assembly stage can be challenging. Therefore, the cloning assembly protocol should be as simple as possible with minimal DNA parts to enhance assembly efficiency. In addition, characterization of the combinatorial library showed population drift and gRNA over-representation, possibly due to PCR amplification bias (Aird et al., 2011). Third, the high DNA concentrations required for large library transformations in yeast resulted in multiple plasmids per cell. Since deconvoluting individual hits to identify the plasmid that increases metabolite production is laborious, alternative strategies such as using CEN plasmids for the gRNA library, which offer better segregation fidelity than 2μ plasmids, could mitigate this issue (Chlebowicz-Śledziewska & Śledziewski, 1985; Murray & Cesareni, 1986).

Taken together, this case study highlights the potential of combining aTF-based biosensors with genome-wide CRISPR screens, providing a powerful tool for targeted genome evolution in a metabolic engineering context. The senMAGIC framework described here enabled us to identify both intuitive and non-intuitive genetic targets for the extensively studied shikimate and ccM pathways, for which multimillion-dollar global markets exist (Averesch & Krömer, 2018), and may prove particularly valuable for engineering less studied pathways or non-conventional yeasts (Pyne et al., 2023; Sun et al., 2020; Thornbury et al., 2025). The success of senMAGIC is in part due to improvements in transformation and gene editing efficiency in *S. cerevisiae* (Gietz & Schiestl, 2007; Horwitz et al., 2015), and could be enhanced when combined with lab automation frameworks (Hillson et al., 2019; Holowko et al., 2021). Furthermore, gene targets identified in this study could be optimized using bidirectional CRISPR systems for precise control of gene expression (Bowman et al., 2020) or through promoter replacement in CRISPR-free strains. In conclusion, senMAGIC establishes biosensor-directed targeted genome evolution as a scalable and generalizable strategy for uncovering new engineering targets in both well-characterized and underexplored metabolic pathways.

## Methods

### Strains and growth media

*Saccharomyces cerevisiae* manipulations were conducted in the prototroph CEN.PK113-7D, which was grown at 30 °C in liquid or solid Yeast Peptone Dextrose medium (YPD; 10 g L^−1^ yeast extract, 20 g L^−1^ peptone, 20 g L^−1^ dextrose) for ∼16 h or 48 h, respectively, unless otherwise stated. Biosensor characterization was performed in Yeast Nitrogen Base medium (YNB; 6.7 g L^−1^ yeast nitrogen base, 20 g L^−1^ dextrose) and Synthetic Complete medium (SC; 6.7 g L^−1^ yeast nitrogen base with ammonium sulfate, 20 g L^−1^ dextrose, 1.92 g L^−1^ yeast synthetic drop-out medium supplement without histidine, 0.08 g L^−1^ histidine) adjusted to pH 4. For MAGIC experiments, modified SC-URA medium (1 g L^−1^ yeast nitrogen base without ammonium sulfate, 1.7 g L^−1^ monosodium glutamate, 20 g L^−1^ dextrose, 1.92 g L^−1^ yeast synthetic drop-out medium supplement without uracil) was used to enable G418 selection. Hygromycin (200 μg ml^−1^) and G418 (200 μg ml^−1^) were added for plasmid selection as needed. Tables S1 and S2 contain a list of strains and plasmids employed in this study, respectively.

### Chemicals

Chemicals for biosensor testing were purchased from Sigma Aldrich and dissolved in water, unless otherwise specified, to obtain the stock concentrations in parentheses: catechol (100 mM), protocatechuate (50 mM), *cis,cis*-muconic acid (5 mM), and adipic acid (100 mM).

### Synthetic genes, oligonucleotides, and plasmids

When needed, synthetic genes were codon-optimized for yeast expression using the IDT tool and purchased from Twist Bioscience. Oligonucleotides were purchased from Thermo Scientific and suspended in nuclease-free water to a concentration of 100 μM. Tables S3 and S4 contain a list of genetic parts and oligonucleotides used in this study, respectively.

### *Saccharomyces cerevisiae* general strain construction

All *S. cerevisiae* genetic modifications were made using CRISPR-Cas9-mediated genomic integration and *in vivo* DNA homologous recombination (Horwitz et al., 2015; Shao et al., 2009). Briefly, yeast cells were transformed with a DNA pool comprising the following parts as needed: PCR-amplified promoters, terminators, and coding sequences with ∼40 bp homology to the preceding and/or following part, mixed in equimolar ratios; two ∼500 bp homology arms to the genomic locus targeted for integration (200 ng each); linearized pCas-G418 and pCas-Hyg vectors encoding Cas9 (100 ng each) (Table S2); PCR-amplified gRNAs targeting the chromosomal locus of choice and with homology to the Cas9 plasmid for assembly of the plasmid by *in vivo* homologous recombination (200 ng per transformation). Unless otherwise stated, the DNA pool was transformed into yeast using a 25 or 50 μL lithium acetate/PEG heat shock transformation (Gietz & Schiestl, 2007) in which cells were incubated at 30 °C for 15 min and heat-shocked at 42 °C for 30 min, followed by recovery for 16 hours in YPD, and plating on YPD-Hyg/G418 dual selection plates. Successful genomic integrations were confirmed by Phire (Thermo Scientific) colony PCR and Sanger or Nanopore amplicon sequencing. Finally, clones were purified and the Cas9 plasmid was cured by streaking colonies two times on YPD plates with no selection with colony PCR confirmations after every step. Sequences of promoters, genes, and terminators are listed in Table S3. In general, Flagfeldt sites 16, 18, and 20 were used for biosensor, enzyme expression, and zeocin resistance cassette integration, respectively (Bourgeois et al., 2018; Flagfeldt et al., 2009). The only exception is the ccM-producing DT64 strains which have the zeocin resistance cassette integrated in Flagfeldt site 14.

For choosing biosensors, preference was given to repressors due to their standardized implementation as genetic circuit components in eukaryotes with respect to the operator sequence position within their regulated promoter compared to activators (Ambri et al., 2020; Murphy et al., 2007). The aTF repressor candidate sensors used in this work included PcaV from *Comamonas testosteroni* (Davis et al., 2013) and Caq101, a variant of the aTF RolR from *Corynebacterium glutamicum* that we previously engineered for responsiveness to catechol (Nasr et al., 2023). Activator-based biosensors included in this work were BenM^WT^ for ccM and BenM^Tisno^ variant responsive to both ccM and adipic acid, derived from *Acinetobacter baylyi* (Skjoedt et al., 2016; Snoek et al., 2020). We integrated the biosensor candidates into the genome of *S. cerevisiae* as single copy genetic circuits that contain both a regulatory and a reporter module as previously described (Nasr et al., 2022; Pham et al., 2024). The regulatory module comprised the candidate biosensor that had its expression driven by the strong constitutive yeast promoter *P_TDH3_*, with different promoters (*P_PGK1_*, *P_CCW12_*, and *P_TEF1_* for repressors, and *P_CYC1_*for activators) driving reporter expression. For the repressor candidates, the reporter promoters were engineered for candidate aTF regulation by inserting a single copy of the operator 11 bp downstream of the terminal TATA box sequence TATAWAWR (W=A/T R=A/G)(Murphy et al., 2007). Given that it is the only activator in our collection of sensor candidates, BenM-regulated promoters were constructed using the previously described and optimized ccM- and/or adipic acid-inducible truncated *P_CYC1_* promoter sequence (Ambri et al., 2020; Skjoedt et al., 2016; Snoek et al., 2020).

### *S. cerevisiae* biosensor characterization

Cells were grown overnight at 30 °C and 300 rpm in 96 deep-well plates in an orbital shaker. The following day, yeast cells were diluted 1:500 into 500 μL SC or YNB medium to an OD_600_ of ∼0.01 in 96 deep-well plates with or without different concentrations of small molecules from the low μM range up to 5-100 mM (depending on the toxicity and solubility of the molecule). The plates were then incubated in an orbital shaker at 30 °C and 300 rpm for 20 h, after which fluorescence was measured using a BD Accuri C6 flow cytometer. Cells were diluted 1:5 in deionized water to obtain an average rate of ∼1000 events/second for a total of 10000 single-cell events. The mean fluorescence of the total ungated population was plotted for each molecule and/or concentration tested. Unless otherwise stated, dose-response data points are averages of triplicates, shown either as 3 data points, or a single data point with an error bar representing standard deviation.

### Zeocin resistance characterization

Cells were grown overnight at 30 °C and 300 rpm in 96 deep-well plates in an orbital shaker. The following day, yeast cells were diluted 1:1000 into 100 μL YPD medium to an OD_600_ of ∼0.005 in 96 well plates with or without different concentrations of zeocin (∼150-300 μg ml^-1^ up to 10-20 mg ml^-1^) and small molecules (∼0.3 mM up to 5-10 mM).

### Construction of senMAGIC platform strains and transformation and selection of pooled gRNA library

senMAGIC yeast strain libraries were constructed by first deleting *URA3* followed by integrating the *PmeI*-digested vector pAID6 containing the three Cas proteins into delta sites and using G418 for selection to generate strains MN3034 and MN3037 (which was further modified by deleting the native *PAD1* to create MN3038) (Table S1). These strains were then transformed with 10 μg CRISPRa, 10 μg CRISPRi, and 20 μg CRISPRd plasmid libraries, into 10 OD_600_ units of strain MN3034, and 40 OD_600_ units of strains MN3037 and MN3038. Transformations were done using the lithium acetate/PEG heat shock transformation (Gietz & Schiestl, 2007) with a modified 1 h heat shock at 42 °C instead of 0.5 h. Cells were then pelleted and suspended in YPD medium (4 ml for MN3034 and 8 ml for MN3037 or MN3038) for a ∼4 h recovery period at 30 °C. Cells were then diluted 1000-fold and spread onto SC-URA/G418 agar plates to evaluate transformation efficiency. Of the remaining cells, 1 ml was pelleted and suspended in 10 mL SC-URA/G418 medium for 48 h. This was followed by zeocin selection for 18 h, for which 0.3 OD_600_ units of strain MN3034 was added to 15 ml SC-URA/G418 with zeocin 900 μg ml^-1^, and 0.6 OD_600_ units of strains MN3037 or MN3038 were added to 15 ml SC-URA/G418 with zeocin 600 μg ml^-1^.

### Preliminary Single gRNA library automated screening workflow

Different dilutions of cells post zeocin-selection were plated on SC-URA/G418 plates and incubated for 48-72 hours to obtain isolated colonies. Then 960 colonies were picked from agar plates using a colony picker (QPix 460) into ten 96 well-plates containing SC-URA/G418 medium. These clones were incubated for 18 h, diluted 1 in 100 to an approximate OD_600_ of 0.01 the following day and incubated for a further 24 h. This was followed by flow cytometry analyses on a BD Accuri C6 flow cytometer. The ∼96 clones with the highest fluorescence were subjected to MS metabolite quantification and Sanger sequencing for gRNA identification.

### gRNA enrichment analysis

Each library sample had 0.3 OD_600_ units harvested by centrifugation and resuspended in 5 μL of 15 % SDS. Samples were boiled in the microwave for 2 minutes and centrifuged at 4000 g for 3 min. All 5 μL of the sample were used as template in a 100 μL Phusion (Thermo Scientific) PCR reaction with primers containing heterogeneity spacers for 20 cycles. A second Phusion PCR was performed using primers with Illumina dual index adapter sequences for 20 cycles. All amplicons were pooled and run on a 0.8 % agarose gel before gel extracting all sequences between 300-500 bp. The purified amplicons were sequenced on an Illumina MiSeq platform using 300 paired-end reads (Genome Quebec). Initial quality control was done using FastQC v0.11.9 and MultiQC v1.12. The paired reads were merged using PEAR v0.9.6 with a minimum read length(-t) of 20 bp and a quality score threshold (-q) of 25. These merged reads were then analyzed for guide sequences using python.

### Applying genome activations, interferences, and deletions to selected genes

For activations and interferences, gRNAs targeting an area proximal to the transcription site of the gene of interest were designed using the CCTop tool (Stemmer et al., 2015). These were combined with either the *P_TDH3_* or a truncated *P_CYC1_* promoters as donor templates for activation and interference, respectively (Table S3), each containing ∼40 bp homology arms with a mutation in the PAM site. For deletions, gRNAs targeting the coding sequence were designed and combined with donor templates encoding 200 bp up- and down-stream of the gene of interest with 40 bp homology to each other, resulting in deletion of the full coding sequence. Activations, interferences, and deletions were confirmed by colony PCR. Sequences of gRNAs used are listed in Table S5.

### LC-MS metabolite analysis

For the quantification of PCA and ccM, cells were grown overnight at 30 °C and 300 rpm in 96 deep-well plates in an orbital shaker. Cells were then diluted to an OD_600_ of 0.1 in either SC-URA/G418 medium for preliminary guide characterization, or Kuhner deep-well plates containing 15 mg ml^-1^ slow release glucose supplemented with SC medium without glucose (starting pH 5.4 for PCA-producing and Tris-HCl-controlled pH 7 for ccM-producing strains) for 24-72 h. Metabolites were extracted directly from the culture broth containing both cells and medium by combining 25 μL of culture with 100 μL of 100% acetonitrile (ACN), shaking at 800 rpm for 5 min, followed by adding 542 μL of sterile ddH_2_O to give a final concentration of 15% (v/v) ACN. More productive samples that lacked pAID6 and the gRNA plasmids (MN3039-MN3068) were diluted a further ∼20 fold. The extracted material was separated on a 1290 Infinity II Liquid chromatography system (Agilent Technologies) using a Zorbax Eclipse Plus C18 column (2.1 x 50 mm; 1.8 µm; Agilent Technologies) and an autosampler tray cooled at 4°C. Metabolites were separated using the following gradient of solvent A (0.1% formic acid in water) and solvent B (0.1% formic acid in 100% ACN): 2% B to 10% B from 0 to 4 min (0.3 mL min^−1^), 10% B to 85% B from 4 to 6 min (0.3 mL min^−1^), held at 85% B from 6 to 7 min (0.3 mL min^−1^), 85% B to 2% B from 7 to 7.1 min (0.3 mL min^−1^), and held at 2% B from 7.1 to 9 min (0.45 mL min^−1^). After separation, the eluent was injected into a 6560 Ion Mobility Q-TOF (Agilent Technologies) using a 100 to 1100 *m/z* scanning range in negative mode. Resolution, capillary voltage, and source temperature were set to 100,000, 5 kV, and 350 °C, respectively. Data was processed using the Agilent Masshunter Quantitative Analysis software.

### Combinatorial library building and screening

(see Supplemental method 1 for details). Briefly, selected gRNAs were amplified from the activation, interference, or deletion genome-wide libraries using one primer specific to the scaffold sequence (A, I, or D) and one primer specific to the gRNA sequence to amplify. Overhangs on the primers contained a BsaI recognition site together with a cleavage site compatible with site 1, 2, or 3 of the gRNA array in the final plasmid. The tRNA_GLY_ splicing unit parts were amplified by the annealing and extension of specific pairs of primers, and the different parts were assembled by Golden Gate using BsaI. The assembly was next purified by ethanol-precipitation and transformed into *E. coli* DH5α before liquid growth and culture saving at -80C. Several transformations were performed to achieve 8.10^5^ and 6.10^5^ transformants for M1 and M3, respectively. Culture vials from -80 °C were next thawed and used to inoculate liquid culture, which were grown until saturation prior to plasmid extraction. M1 and M3 were next transformed with 200 µg of plasmids with the same protocol as for the single-gRNA transformation. Libraries were next inoculated at OD=0.05 in triplicates in 25ml SC-URA with G418 and zeocin (6.5 mg mL^−1^ and 24 h for M1, and 8 mg mL^−1^ and 48 h for M3). Cells were next diluted at OD=0.2 in SC-URA with G418 and zeocin 1 mg mL^−1^ and grown for 4 h for M1 and 24 h for M3 prior to FACS sorting (FACSMelody, BD). Cells with the highest GFP signal were sorted, recovered, and plated. Single colonies were next inoculated into a regular 96 well plate for 16 hours in SC-URA with G418 and diluted at OD=0.2 in 96-deepwell plates in the same media for 24 hours growth before metabolite quantification. The top PCA or ccM producers were Sanger-sequenced for gRNA identification.

### Quality control of the cloned combinatorial library

(see Supplemental method 2 for details). Briefly, pooled bacterial cultures from the multiple cloning rounds were spread on solid selective LB media to obtain single colonies. 96 colonies for each library (M1 and M3) were then resuspended in 50 µL deionized water in two 96-well format PCR plates using a colony picker (QPix, VWR) and the cell suspensions were boiled for 2 minutes in a thermocycler. Boiled suspended colonies, PCR master mix, and barcoded primers were next dispensed in a 384-well format PCR plate with an acoustic liquid handler (Echo, Beckman Coulter) in a total of a 6µL reaction. The 2x96 PCR reactions were next pooled together, gel-purified, and sequenced with a Nanopore device (Oxford Nanopore). Reads were next demultiplexed and analyzed for guides identification.

## CRediT authorship contribution statement

**Mohamed A. Nasr:** Conceptualization, Investigation, Methodology, Formal analysis, Visualisation, Writing – Original Draft Preparation, Writing – Review & Editing. **Adrien Knoops:** Conceptualization, Investigation, Methodology, Formal analysis, Visualisation, Writing – Review & Editing. **Mackenzie Thornbury:** Conceptualization, Investigation, Methodology, Formal analysis, Visualisation, Writing – Review & Editing. **Meghan Davies:** Conceptualization, Investigation, Methodology, Formal analysis, Visualisation. **David H. Kwan:** Conceptualization, Funding Acquisition, Supervision, Writing – Review & Editing. **Vincent J.J. Martin:** Conceptualization, Funding Acquisition, Supervision, Writing – Review & Editing

## Acknowledgments

This study was supported by NSERC through discovery grants to V.J.J.M and D.H.K. M.A.N. was supported by a Concordia International Tuition Award of Excellence, and an FRQNT B2X Doctoral Research Award. A.K. was supported by a Concordia Horizon Postdoctoral Fellowship. M.T. was supported by a FRQNT B2X Doctoral Research Award, a SynBioApps NSERC-CREATE Scholarship, and an NSERC Vanier Canada Graduate Scholarship. V.J.J.M. was supported by a Concordia University Research Chair. We thank Dr. Malcolm Whiteway for critical reading of the manuscript. We thank Dr. Radhakrishnan Mahadevan for providing lab resources to construct strains in figure 5. We thank Dr. Huimin Zhao for kindly providing the gRNA libraries used in this work.

## Data availability

Data will be made available on request.

