## Supplementary information for "Biosensor-guided genome-wide CRISPR screens enable *cis*,*cis*-muconic acid overproduction in yeast"

Footnotes:

1 Present Address: BioZone, Department of Chemical Engineering and Applied Chemistry, University of Toronto, Toronto, Ontario, Canada

2 Present Address: Institut des Science du vivant, HES-SO, Rue de l’industrie 19, 1950 Sion, VS, Switzerland

3 Present Address: pHathom technologies, Halifax, Nova Scotia, Canada

4 Present Address: BenchSci, Toronto, Ontario, Canada

**Supplementary information**

**Table S1. List of strains used in this work.**

| **Strain ID** | **Parent** | **Genotype or integrated cassette** | **Locus** | | **Source** |
| --- | --- | --- | --- | --- | --- |
| CEN.PK113-7D | CEN.PK113-7D | Genotype: *MATα MAL2-8c SUC2* | N.A | (Nijkamp et al., 2012) | |
| LB329 | CEN.PK113-13D | *ARO3*Δ *ARO4*Δ | N.A | (Pyne et al., 2018) | |
| DT64 | CEN.PK113-13D | P*_TDH_*_3_-*AROZ*- T*_TDH2_*- P*_TEF1_*-*aroY*- T*_ADH2_*  P*_TEF1_*-*ARO4^fbr^*- T*_ADH2_*- P*_TDH_*_3_-*ARO3^fbr^*-T*_TDH1_*  P*_TDH_*_3_-*HQD2*-T*_TDH1_*  P*_TDH_*_3_-*HQD2*- T*_ADH1_*  P*_PGK1_*-*PAD1*- T*_ADH2_*  P*_PGK1_*-*aroB*-T*_ADH1_*- P*_TDH_*_3_-*aroD*-T*_Eno1_*-*URA3* | FgF18  FgF19  FgF20  USERXII-5  USERXII-2  URA | Daniel Tsyplenkov, unpublished | |
| MN3001 | CEN.PK113-7D | P*_TDH_*_3_-*caq101*-T*_Eno2_*-P*_CCW12_*-*gfp^Envy^*-T*_ADH1_* | FgF16 | This work | |
| MN3004 | CEN.PK113-7D | P*_TDH_*_3_-*pcaV*-T*_Eno2_*-P*_CCW12_*-*gfp^Envy^*-T*_ADH1_* | FgF16 | This work | |
| Caq101 | CEN.PK113-7D | P*_TDH_*_3_-*caq101*-T*_Eno2_*-P*_CCW12O_*-*gfp^Envy^*-T*_ADH1_* | FgF16 | (Nasr et al., 2023) | |
| MN3009 | CEN.PK113-7D | P*_TDH_*_3_-*pcaV*-T*_Eno2_*-P*_CCW12O_*-*gfp^Envy^*-T*_ADH1_* | FgF16 | This work | |
| MN3012 | CEN.PK113-7D | P*_TDH_*_3_-*benM*-T*_Eno2_*-P*_CYC1O_*-*gfp^Envy^*-T*_ADH1_* | FgF16 | This work | |
| MN3013 | CEN.PK113-7D | P*_TDH_*_3_-*benM^Tisno^-*T*_Eno2_*-P*_CYC1O_*-*gfp^Envy^*-T*_ADH1_* | FgF16 | This work | |
| MN3013 | CEN.PK113-7D | P*_TDH_*_3_-*caq101*-T*_Eno2_*-P*_TEF1_*-*gfp^Envy^*-T*_ADH1_* | FgF16 | This work | |
| MN3014 | CEN.PK113-7D | P*_TDH_*_3_-*caq101*-T*_Eno2_*-P*_TEF1O_*-*gfp^Envy^*-T*_ADH1_* | FgF16 | This work | |
| MN3015 | CEN.PK113-7D | P*_TDH_*_3_-*caq101*-T*_Eno2_*-P*_PGK1_*-*gfp^Envy^*-T*_ADH1_* | FgF16 | This work | |
| MN3016 | CEN.PK113-7D | P*_TDH_*_3_-*caq101*-T*_Eno2_*-P*_PGK1O_*-*gfp^Envy^*-T*_ADH1_* | FgF16 | This work | |
| MN3017 | CEN.PK113-7D | P*_TDH_*_3_-*pcaV*-T*_Eno2_*-P*_TEF1_*-*gfp^Envy^*-T*_ADH1_* | FgF16 | This work | |
| MN3018 | CEN.PK113-7D | P*_TDH_*_3_-*pcaV*-T*_Eno2_*-P*_TEF1O_*-*gfp^Envy^*-T*_ADH1_* | FgF16 | This work | |
| MN3019 | CEN.PK113-7D | P*_TDH_*_3_-*pcaV*-T*_Eno2_*-P*_PGK1_*-*gfp^Envy^*-T*_ADH1_* | FgF16 | This work | |
| MN3020 | CEN.PK113-7D | P*_TDH_*_3_-*pcaV*-T*_Eno2_*-P*_PGK1O_*-*gfp^Envy^*-T*_ADH1_* | FgF16 | This work | |
| MN3021 | CEN.PK113-7D | P*_TDH_*_3_-*pcaV*-T*_Eno2_*-P*_CCW12OO_*-*gfp^Envy^*-T*_ADH1_* | FgF16 | This work | |
| MN3022 | CEN.PK113-7D | P*_TDH_*_3_-*pcaV*-T*_Eno2_*-P*_TEF1OO_*-*gfp^Envy^*-T*_ADH1_* | FgF16 | This work | |
| MN3023 | CEN.PK113-7D | P*_TDH_*_3_-*pcaV*-T*_Eno2_*-P*_PGK1OO_*-*gfp^Envy^*-T*_ADH1_* | FgF16 | This work | |
| MN3024 | CEN.PK113-7D | P*_TDH_*_3_-*pcaV*-T*_Eno2_*-P*_CCW12OO_*-*gfp^Envy^*-T*_ADH1_*  P*_PGK1_*-*AROZ*-T*_ENO2_* | FgF16  FgF18 | This work | |
| MN3025 | CEN.PK113-7D | P*_TDH_*_3_-*pcaV*-T*_Eno2_*-P*_CCW12OO_*-*gfp^Envy^*-T*_ADH1_*  P*_PGK1_*-*AROZ*-T*_ENO2_*  P*_CCW12OO_*-*bleoR*-T*_ADH1_* | FgF16  FgF18  FgF20 | This work | |
| MN3026 | CEN.PK113-7D | P*_TDH_*_3_-*pcaV*-T*_Eno2_*-P*_CCW12OO_*-*gfp^Envy^*-T*_ADH1_* | FgF16 | This work | |
| MN3027 | LB329 | P*_TDH_*_3_-*pcaV*-T*_Eno2_*-P*_CCW12OO_*-*gfp^Envy^*-T*_ADH1_*  P*_PGK1_*-*AROZ*-T*_ENO2_* | FgF16  FgF18 | This work | |
| MN3028 | LB329 | P*_TDH_*_3_-*pcaV*-T*_Eno2_*-P*_CCW12OO_*-*gfp^Envy^*-T*_ADH1_*  P*_PGK1_*-*AROZ*-T*_ENO2_*  P*_CCW12OO_*-*bleoR*-T*_ADH1_* | FgF16  FgF18  FgF20 | This work | |
| MN3029 | CEN.PK113-7D | P*_TDH_*_3_-*caq101*-T*_Eno2_*-P*_CCW12O_*-*gfp^Envy^*-T*_ADH1_*  P*_CCW12O_*-*bleoR*-T*_ADH1_* | FgF16  FgF20 | This work | |
| MN3030 | CEN.PK113-7D | P*_TDH_*_3_-*benM^Tisno^-*T*_Eno2_*-P*_CYC1O_*-*gfp^Envy^*-T*_ADH1_*  P*_CYC1O_*-*bleoR*-T*_ADH1_* | FgF16  FgF14 | This work | |
| MN3031 | DT64 | P*_TDH_*_3_-*benM^Tisno^-*T*_Eno2_*-P*_CYC1O_*-*gfp^Envy^*-T*_ADH1_* | FgF16 | This work | |
| MN3032 | DT64 | P*_TDH_*_3_-*benM^Tisno^-*T*_Eno2_*-P*_CYC1O_*-*gfp^Envy^*-T*_ADH1_*  P*_CYC1O_*-*bleoR*-T*_ADH1_* | FgF16  FgF14 | This work | |
| MN3033 | MN3025 | P*_TDH_*_3_-*pcaV*-T*_Eno2_*-P*_CCW12OO_*-*gfp^Envy^*-T*_ADH1_*  P*_PGK1_*-*AROZ*-T*_ENO2_*  P*_CCW12OO_*-*bleoR*-T*_ADH1_*  *URA3*Δ | FgF16  FgF18  FgF20 | This work | |
| MN3034 (MAGIC 1) | MN3033 | P*_TDH_*_3_-*pcaV*-T*_Eno2_*-P*_CCW12OO_*-*gfp^Envy^*-T*_ADH1_*  P*_PGK1_*-*AROZ*-T*_ENO2_*  P*_CCW12OO_*-*bleoR*-T*_ADH1_*  *URA3*Δ  pAID6 | FgF16  FgF18  FgF20  URA3  Delta sites |  | |
| MN3036 | MN3032 | P*_TDH_*_3_-*benM^Tisno^-*T*_Eno2_*-P*_CYC1O_*-*gfp^Envy^*-T*_ADH1_*  P*_CYC1O_*-*bleoR*-T*_ADH1_*  *URA3*Δ | FgF16  FgF14 | This work | |
| MN3037 (MAGIC3) | MN3036 | P*_TDH_*_3_-*benM^Tisno^-*T*_Eno2_*-P*_CYC1O_*-*gfp^Envy^*-T*_ADH1_*  P*_CYC1O_*-*bleoR*-T*_ADH1_*  *URA3*Δ  pAID6 integrated | FgF16  FgF14 |  | |
| MN3038  (MAGIC3.1) | MN3037 | P*_TDH_*_3_-*benM^Tisno^-*T*_Eno2_*-P*_CYC1O_*-*gfp^Envy^*-T*_ADH1_*  P*_CYC1O_*-*bleoR*-T*_ADH1_*  *URA3*Δ, *PAD1*Δ | FgF16  FgF14 | This work | |
| MN3049 | MN3024 | *P_CYC1_-ARO2* |  | This work | |
| MN3050 | DT64 | *P_CYC1_-ARO2* |  | This work | |
| AK082 | MN3034 | *P_CYC1_-mScarlet-T_ADH1_* | XII-1 | This work | |
| AK122 | MN3031 | P*_TDH_*_3_-*benM^Tisno^-*T*_Eno2_*-P*_CYC1O_*-*gfp^Envy^*-T*_ADH1_*  P*_CYC1O_*-*bleoR*-T*_PGK1_* | FgF16  FgF14 | This work | |
| AK128 | AK122 | P*_TDH_*_3_-*benM^Tisno^-*T*_Eno2_*-P*_CYC1O_*-*gfp^Envy^*-T*_ADH1_*  P*_CYC1O_*-*bleoR*-T*_PGK1_*  *△URA3* | FgF16  FgF14  URA3 | This work | |
| AK180 | AK128 | P*_TDH_*_3_-*benM^Tisno^-*T*_Eno2_*-P*_CYC1O_*-*gfp^Envy^*-T*_ADH1_*  P*_CYC1O_*-*bleoR*-T*_PGK1_*  *△URA3*  *△PAD1* | FgF16  FgF14  URA3  PAD1 | This work | |
| AK181 | AK180 | P*_TDH_*_3_-*benM^Tisno^-*T*_Eno2_*-P*_CYC1O_*-*gfp^Envy^*-T*_ADH1_*  P*_CYC1O_*-*bleoR*-T*_PGK1_*  *△URA3*  *△PAD1*  pAID6 integrated | FgF16  FgF14  URA3  PAD1  Delta sites | This work | |
| MN3051 | MN3049 | *P_CYC1_-IDP1* |  | This work | |
| MN3052 | MN3049 | *P_CYC1_-RVS167* |  | This work | |
| MN3053 | MN3049 | *P_CYC1_-RIM4* |  | This work | |
| MN3054 | MN3049 | *P_CYC1_-TPI1* |  | This work | |
| MN3055 | MN3049 | *P_TDH3_-CCS1* |  | This work | |
| MN3056 | MN3049 | *P_TDH3_-PDR3* |  | This work | |
| MN3057 | MN3049 | *P_CYC1_-IDP1 + P_CYC1_-TPI1* |  | This work | |
| MN3058 | MN3049 | *P_TDH3_-CCS1 + P_TDH3_-PDR3* |  | This work | |
| MN3059 | MN3049 | *P_TDH3_-CCS1 + P_TDH3_-PDR3 + P_CYC1_-RVS167* |  | This work | |
| MN3060 | DT64 | *P_CYC1_-IDP1* |  | This work | |
| MN3061 | DT64 | *P_CYC1_-RVS167* |  | This work | |
| MN3062 | DT64 | *P_CYC1_-RIM4* |  | This work | |
| MN3063 | DT64 | *P_CYC1_-TPI1* |  | This work | |
| MN3064 | DT64 | *P_TDH3_-CCS1* |  | This work | |
| MN3065 | DT64 | *P_TDH3_-PDR3* |  | This work | |
| MN3066 | DT64 | *P_CYC1_-IDP1 + P_CYC1_-TPI1* |  | This work | |
| MN3067 | DT64 | *P_TDH3_-CCS1 + P_TDH3_-PDR3* |  | This work | |
| MN3068 | DT64 | *P_TDH3_-CCS1 + P_TDH3_-PDR3 + P_CYC1_-RVS167* |  | This work | |
| MN3069 | MN3050 | *P_CYC1_-IDP1* |  | This work | |
| MN3070 | MN3050 | *P_CYC1_-RVS167* |  | This work | |
| MN3071 | MN3050 | *P_CYC1_-RIM4* |  | This work | |
| MN3072 | MN3050 | *P_CYC1_-TPI1* |  | This work | |
| MN3073 | MN3050 | *P_TDH3_-CCS1* |  | This work | |
| MN3074 | MN3050 | *P_TDH3_-PDR3* |  | This work | |
| MN3075 | MN3050 | *P_CYC1_-IDP1 + P_CYC1_-TPI1* |  | This work | |
| MN3076 | MN3050 | *P_TDH3_-CCS1 + P_TDH3_-PDR3* |  | This work | |
| MN3077 | MN3050 | *P_TDH3_-CCS1 + P_TDH3_-PDR3 + P_CYC1_-RVS167* |  | This work | |

**Table S2. List of plasmids used in this work**

| **Name** | **Description** |  | **Source** |
| --- | --- | --- | --- |
| pBBK94 | pCas-G418 |  | (Bean et al., 2022) |
| pBBK95 | pCas-Hyg |  | (Bean et al., 2022) |
| pAID6 | Yeast integration plasmid for expression of dLbCpf1-VP, dSpCas9-RD1152, SaCas9, and Csy4 |  | (Lian et al., 2019) |
| p426*-SaSgH | (Empty Backbone) Helper plasmid for SaCas9 gRNA cloning and expression |  | (Lian et al., 2019) |
| p426*-SpSgH | (Empty Backbone) Helper plasmid for SpCas9 gRNA cloning and expression |  | (Lian et al., 2019) |
| p426*-SaSgH-MNmod | Modified empty backbone for combinatorial gRNA assembly |  | This work |
| pGG1 | p426*-SaSgH-MNmod containing guides *CYC1a -CCW12i(wrong design)-ADE2d(wrong design)* |  | This work |
| pAK091 | p426*-SaSgH-MNmod containing guides *CYC1a-CCW12i-ADE2d* |  | This work |
| pAK142 | p426*-SaSgh-MNmod containing a CYC1pa guide |  | This work |
| pAK143 | p426*-SaSgh-MNmod containing a CCW12i guide |  | This work |
| pAK144 | p426*-SaSgh-MNmod containing a ADE2d guide |  | This work |
| pBBK04 | pCas plasmid from Mylo toolkit with SplitHygR system |  | (Bean et al., 2022) |
| pAK215 | *pBBK04::gRNA for FgF14 cut* |  | This work |
| pMT293 | SOE1_i single guide plasmid |  | This work |
| pMT294 | RVS167_i single guide plasmid |  | This work |
| pMT295 | DUG3_i single guide plasmid |  | This work |
| pMT296 | RIM4_i single guide plasmid |  | This work |
| pMT297 | PDR3_a single guide plasmid |  | This work |
| pMT298 | CCS1_a single guide plasmid |  | This work |
| pMT299 | RNH202_a single guide plasmid |  | This work |
| pMT300 | PDR3_a + RIM4_i double guide plasmid |  | This work |
| pMT301 | PDR3_a + DUG3_i double guide plasmid |  | This work |
| pMT302 | PDR3_a + RNH202_a double guide plasmid |  | This work |
| pMT303 | RNH202_a + DUG3_i double guide plasmid |  | This work |
| pMT304 | PDR3_a + CCS1_a double guide plasmid |  | This work |
| pMT305 | CCS1_a + RIM4_i double guide plasmid |  | This work |
| pMT389 | IDP1_i single guide plasmid |  | This work |
| pMT390 | TPI1_i single guide plasmid |  | This work |
| pMT391 | IDP1_i + TPI1_i double guide plasmid |  | This work |
| pMT392 | TPI1_i + IDP1_i double guide plasmid |  | This work |

**Table S3. List of genes and parts used in this work:**

| **Gene ID** | **Sequence** | **Source** |
| --- | --- | --- |
| *caq101* | atgcctacaccctcccagcataaggacgcctccacagcgcaaacagataatcaggtccccactggacgtagggcgcagaagagagagcaaacaagggcaaggttgattacgtcagcgagaacattgatggctgagcgtggcgtggataacgttggaatcgccgaaataacggagggggccaacatcggaaccggtacgttctataattattttcccgatagagaacaattactacaggctgttgcggaagacgcttttgagagtgttggcatagcactagatcaagtacttacaaagttggacgatcccgcggaagtctttgcgggcgggttaagacatcttgtaagacactccctagaagatagaatttgggggggttttttcatccaaatgggggccgcccatccggtcttaatgaggatatttggtccaagagctaggcgtgatttattgcatggtctggagaccggtcgtttcactattgaggatttggacctagccacaacctgcacctttggcagcctaatagctgctatacagatggcattatcagcagatcaggacagtaacgacgacaaagaccaaatttttgcggcagcgatgctgagaatggtaggggtacaagcagcagaagcgagggaaattgcgtctcgtcccctacccgagatcagcccggttaagccgcagtag | (Nasr et al., 2023) |
| *pcaV* | atggcagcagtagaccttgccacgcaccccggtcacttggctagaagattgcaacaagcgcattatttgttatggaacaccatggtaagtgaggagacaacgagtccacagtacgcagtgttgaacgctctggtggcggagccaggtcttgatcaaaggactgtcggcgaacgtgttgggctggatagatctaccatagctgaagtcgtttcaaggcttggcaggaggggtttactagacaaagtaagagaccctcaggatggtaggcgtagcctgcttaggctgacggatgagggtctaagggtacacagaaggctgggagtgaggatcgccagaatgaatcaggtcttcttagccccattggctgcagatgaacaagccgttttcttcgatttgattcgtagagtggcggatgcagccgaaggacttcgtaatccggcagaaccagccgtggctcctggttga | gBlock |
| *benMWT* | atggaattgagacacttgagatacttcgttgccgttgttgaagaacaatcttttacaaaggctgcagacaagttgtgtattgctcaaccaccattatccagacaaatccaaaacttggaagaagaattgggtatccaattattagaaagaggttccagaccagttaagactactccagaaggtcatttcttttaccaatacgccatcaagttgttgtccaacgttgatcaaatggtcagtatgaccaagagaattgcctctgttgaaaagaccattagaatcggttttgttggttccttgttgttcggtttgttgccaagaattatccacttgtacagacaagctcatccaaacttgagaatcgaattatacgaaatgggtactaaggctcaaaccgaagctttgaaagaaggtagaattgacgctggttttggtagattgaagatttctgatccagccatcaagagaaccttgttgagaaacgaaagattgatggttgctgttcatgcttcccatccattgaatcaaatgaaggataagggtgttcacttgaacgatttgatcgacgaaaagatcttgttgtacccatcttctccaaagccaaacttctctactcatgttatgaacatcttctctgaccatggtttggaacctaccaagattaacgaagttagagaagtccaattggccttgggtttggttgctgctggtgaaggtatttcattggttccagcttctacccaatccattcaattattcaacttgtcctacgtcccattattagatccagatgctattaccccaatctacattgctgttagaaacatggaagaatccacctacatctactcattatacgaaaccatcagacaaatctacgcctacgaaggttttactgaaccaccaaattggtaa | gBlock |
| *benMTisno* | atggaattgagacacttgagatacttcgttgccgttgttgaagaacaatcttttacaaaggctgccgacaagttgtgtattgctcaaccaccattatccagacaaatccaaaacttggaagaagaattgggtatccaattattggaaagaggttccagaccagttaagactactccagaaggtcatttcttttaccaatacgccatcaagttgttgtccaacgttgatcaaatggtcagtatgaccaagagaattgcctctgttgaaaagaccattagaatcggttttgttggttccttgttgttcggtttgttgccaagaattatccacttgtacagacaagctcatccaaacttgagaatcgaattatacgaaatgggtactaaggatcaaaccgaagctttgaaagaaggtagaattgacgctggttttggtagattgaagatttctgatccaggtatcaagagaaccttgttgagaaacgaaagattgatggttgctgttcatgcttcccatccattgaatcaaatgaaggataagggtgttcacttgaacgatttgatcgacgaaaagatcttgttgtacccatcttctccaaagtctaacttctctactcatgttatgaacatcttctctgaccatggtttggaacctaccaagattaacgaagttagagaagtccaattggccttgggtttggttgctgctggtgaaggtatttcattggttccagcttctacccaatccattcaattattcaacttgtcctacgtcccattattagatccagatgctattaccccaatctacattgctgttagaaacatggaagaatccacctacatctactcattatacgttaccatcagacaaatctacgcctacgaaggttttactgaaccaccaaattggtaa | gBlock |
| *gfpEnvy* | atgtctaaaggcgaggaattgtttacaggtgtagttcctattttggtcgaactagatggagatgtcaacggtcataagttttctgtacgtggtgaaggcgaaggtgatgctactaatggcaaactgactctgaagttcatctgtacgaccggaaaattgccagttccatggcctacattagtaacaacattatgttacggagttcaatgctttgctagatacccagatcacatgaagcagcatgactttttcaaaagtgctatgccagaaggttatgtgcaggagagaactatctccttcaaagatgatggtacttataagacaagagcagaggtaaagtttgaaggcgatacattggttaatagaattgagcttaaaggcattgattttaaggaagatgggaatatcctcggacataaacttgaatacaatttcaattcacataatgtttatatcacagcagataaacaaaagaatggtataaaagcgaactttaagacaagacataacgtggaagatggtagcgtccaattagccgaccattaccaacaaaacactcctatcggggacgggccagttctccttccagacaaccactacctatcaacccaatctgttctatctaaagaccctaatgaaaaacgagatcacatggtcttgttagagttcgtgactgccgctggtataacattaggaatggacgaattgtacaaataa | gBlock |
| *AROZ* | *atgccatccaagttggccatcacctctatgtctttgggtagatgttatgctggtcattctttcactaccaagttggatatggctagaaagtatggttaccaaggtttggaattattccacgaagatttggctgatgttgcctatagattgtctggtgaaactccatctccatgtggtccatcaccagctgctcaattgtctgctgctagacaaattttgagaatgtgccaagtcagaaacatcgaaatcgtttgcttgcaaccattctctcaatacgatggtttgttggatagagaagaacacgaaagaagattggaacaattggaattctggatcgaattggcccatgaattggataccgatattattcaaattccagccaacttcttgccagccgaagaagttactgaagatatctctttgattgtctccgacttgcaagaagtagctgatatgggtttacaagctaacccaccaattagattcgtttacgaagctttgtgttggtccactagagttgatacttgggaaagatcttgggaagttgttcaaagagttaacagaccaaacttcggtgtttgtttggacacttttaacattgccggtagagtttatgctgatccaactgttgcttctggtagaactccaaatgctgaagaagctatcagaaagtccattgccagattggttgaaagagttgacgtttccaaggttttctacgttcaagttgttgatgccgaaaagttgaagaaaccattggttccaggtcacagattctatgatccagaacaaccagctagaatgtcttggtctagaaactgcagattattctacggtgaaaaggatagaggtgcttacttgccagtaaaagaaattgcttgggcttttttcaacggtttgggttttgaaggttgggtttccttagaattattcaatagaagaatgtccgacaccggttttggtgttccagaagaattagctagaagaggtgctgtttcttgggctaaattggttagagatatgaagatcaccgttgactctccaactcaacaacaagctacacaacaacctatcagaatgttgtctttatccgctgccttgtaa* | DT64 |
| *bleoR* | *atggctaaattaacatctgccgttcctgttttaacagctagggatgttgcaggtgctgtagagttttggacagataggttaggattctcaagagactttgttgaggacgattttgctggtgttgtcagggatgacgttactttatttatctcagcagtccaagatcaagttgtccctgataatacattggcttgggtctgggtcaggggtttagatgaattatatgctgaatggtcagaagttgtatctacaaacttcagagatgcttctggtccagctatgaccgagattggtgaacagccatggggtagagaatttgctttgagagatccagctggaaattgtgttcattttgttgctgaagaacaagattaa* | pYTK080 |
| **Promoter ID** | **sequence** |  |
| *TDH3* | tcgagtttatcattatcaatactgccatttcaaagaatacgtaaataattaatagtagtgattttcctaactttatttagtcaaaaaattagccttttaattctgctgtaacccgtacatgcccaaaatagggggcgggttacacagaatatataacatcgtaggtgtctgggtgaacagtttattcctggcatccactaaatataatggagcccgctttttaagctggcatccagaaaaaaaaagaatcccagcaccaaaatattgttttcttcaccaaccatcagttcataggtccattctcttagcgcaactacagagaacaggggcacaaacaggcaaaaaacgggcacaacctcaatggagtgatgcaacctgcctggagtaaatgatgacacaaggcaattgacccacgcatgtatctatctcattttcttacaccttctattaccttctgctctctctgatttggaaaaagctgaaaaaaaaggttgaaaccagttccctgaaattattcccctacttgactaataagtatataaagacggtaggtattgattgtaattctgtaaatctatttcttaaacttcttaaattctacttttatagttagtcttttttttagttttaaaacaccaagaacttagtttcgaaaaaca | *S. cerevisiae* genomic DNA |
| *CCW12* | cacccatgaaccacacggttagtccaaaaggggcagttcagattccagatgcgggaattagcttgctgccaccctcacctcactaacgctgcggtgtgcggatacttcatgctatttatagacgcgcgtgtcggaatcagcacgcgcaagaaccaaatgggaaaatcggaatgggtccagaactgctttgagtgctggctattggcgtctgatttccgttttgggaatcctttgccgcgcgcccctctcaaaactccgcacaagtcccagaaagcgggaaagaaataaaacgccaccaaaaaaaaaaaaataaaagccaatcctcgaagcgtgggtggtaggccctggattatcccgtacaagtatttctcaggagtaaaaaaaccgtttgttttggaatttcccatttcgcggccacctacgccgctatctttgcaacaactatctgcgataactcagcaaattttgcatattcgtgttgcagtattgcgataatgggagtcttacttccaacataacggcagaaagaaatgtgagaaaattttgcatcctttgcctccgttcaagtatataaagtcggcatgcttgataatctttctttccatcctacattgttctaattattcttattctcctttattctttcctaacataccaagaaattaatcttctgtcattcgcttaaacactatatcaataa | *S. cerevisiae* genomic DNA |
| *TEF1* | cttcatcggtatcttcgctatattctttttagtcgaatttgcggggagaagatggatctatgctaaactaaataggcatttgaaaaacgacgacgagttacacgacatatcgccatctttaaatgagcaaccacactgggacctcatagaggacgggtctcgctggagtaaatttttcaacgggataattaagacgacaagaaggttcacgaaatctttaatgaggtctttagtcagaggcaggaacagccgtcaagggggcataagactacggtcatccccatctgcctcttcgtccagccttgccaacagggagttcttcagagacatggaggctcaaaacgaaattattgacagcctagacatcaatagtcatacaacagaaagcgaccacccaactttggctgataatagcgtataaacaatgcatactttgtacgttcaaaatacaatgcagtagatatatttatgcatattacatataatacatatcacataggaagcaacaggcgcgttggacttttaattttcgaggaccgcgaatccttacatcacacccaatcccccacaagtgatcccccacacaccatagcttcaaaatgtttctactccttttttactcttccagattttctcggactccgcgcatcgccgtaccacttcaaaacacccaagcacagcatactaaatttcccctctttcttcctctagggtgtcgttaattacccgtactaaaggtttggaaaagaaaaaagagaccgcctcgtttctttttcttcgtcgaaaaaggcaataaaaatttttatcacgtttctttttcttgaaaatttttttttttgatttttttctctttcgatgacctcccattgatatttaagttaataaacggtcttcaatttctcaagtttcagtttcatttttcttgttctattacaactttttttacttcttgctcattagaaagaaagcatagcaatctaatctaagttttaattacaaaaaaaca | *S. cerevisiae* genomic DNA |
| *PGK1* | acgcacagatattataacatctgcataataggcatttgcaagaattactcgtgagtaaggaaagagtgaggaactatcgcatacctgcatttaaagatgccgatttgggcgcgaatcctttattttggcttcaccctcatactattatcagggccagaaaaaggaagtgtttccctccttcttgaattgatgttaccctcataaagcacgtggcctcttatcgagaaagaaattaccgtcgctcgtgatttgtttgcaaaaagaacaaaactgaaaaaacccagacacgctcgacttcctgtcttcctattgattgcagcttccaatttcgtcacacaacaaggtcctagcgacggctcacaggttttgtaacaagcaatcgaaggttctggaatggcgggaaagggtttagtaccacatgctatgatgcccactgtgatctccagagcaaagttcgttcgatcgtactgttactctctctctttcaaacagaattgtccgaatcgtgtgacaacaacagcctgttctcacacactcttttcttctaaccaagggggtggtttagtttagtagaacctcgtgaaacttacatttacatatatataaacttgcataaattggtcaatgcaagaaatacatatttggtcttttctaattcgtagtttttcaagttcttagatgctttctttttctcttttttacagatcatcaaggaagtaattatctactttttacaacaaatataaaaca | *S. cerevisiae* genomic DNA |
| *CCW12-rolo* | cacccatgaaccacacggttagtccaaaaggggcagttcagattccagatgcgggaattagcttgctgccaccctcacctcactaacgctgcggtgtgcggatacttcatgctatttatagacgcgcgtgtcggaatcagcacgcgcaagaaccaaatgggaaaatcggaatgggtccagaactgctttgagtgctggctattggcgtctgatttccgttttgggaatcctttgccgcgcgcccctctcaaaactccgcacaagtcccagaaagcgggaaagaaataaaacgccaccaaaaaaaaaaaaataaaagccaatcctcgaagcgtgggtggtaggccctggattatcccgtacaagtatttctcaggagtaaaaaaaccgtttgttttggaatttcccatttcgcggccacctacgccgctatctttgcaacaactatctgcgataactcagcaaattttgcatattcgtgttgcagtattgcgataatgggagtcttacttccaacataacggcagaaagaaatgtgagaaaattttgcatcctttgcctccgttcaagtatataaagtcggcatgcaaaatctgaacccttgttcatttatgaatcatgattcattcttattctcctttattctttcctaacataccaagaaattaatcttctgtcattcgcttaaacactatatcaataa | *S. cerevisiae* genomic DNA |
| *PGK1-rolo* | acgcacagatattataacatctgcataataggcatttgcaagaattactcgtgagtaaggaaagagtgaggaactatcgcatacctgcatttaaagatgccgatttgggcgcgaatcctttattttggcttcaccctcatactattatcagggccagaaaaaggaagtgtttccctccttcttgaattgatgttaccctcataaagcacgtggcctcttatcgagaaagaaattaccgtcgctcgtgatttgtttgcaaaaagaacaaaactgaaaaaacccagacacgctcgacttcctgtcttcctattgattgcagcttccaatttcgtcacacaacaaggtcctagcgacggctcacaggttttgtaacaagcaatcgaaggttctggaatggcgggaaagggtttagtaccacatgctatgatgcccactgtgatctccagagcaaagttcgttcgatcgtactgttactctctctctttcaaacagaattgtccgaatcgtgtgacaacaacagcctgttctcacacactcttttcttctaaccaagggggtggtttagtttagtagaacctcgtgaaacttacatttacatatatataaacttgcataaaaaaatctgaacccttgttcatttatgaatcatgattcattcgtagtttttcaagttcttagatgctttctttttctcttttttacagatcatcaaggaagtaattatctactttttacaacaaatataaaaca | *S. cerevisiae* genomic DNA |
| *TEF1-rolo* | cttcatcggtatcttcgctatattctttttagtcgaatttgcggggagaagatggatctatgctaaactaaataggcatttgaaaaacgacgacgagttacacgacatatcgccatctttaaatgagcaaccacactgggacctcatagaggacgggtctcgctggagtaaatttttcaacgggataattaagacgacaagaaggttcacgaaatctttaatgaggtctttagtcagaggcaggaacagccgtcaagggggcataagactacggtcatccccatctgcctcttcgtccagccttgccaacagggagttcttcagagacatggaggctcaaaacgaaattattgacagcctagacatcaatagtcatacaacagaaagcgaccacccaactttggctgataatagcgtataaacaatgcatactttgtacgttcaaaatacaatgcagtagatatatttatgcatattacatataatacatatcacataggaagcaacaggcgcgttggacttttaattttcgaggaccgcgaatccttacatcacacccaatcccccacaagtgatcccccacacaccatagcttcaaaatgtttctactccttttttactcttccagattttctcggactccgcgcatcgccgtaccacttcaaaacacccaagcacagcatactaaatttcccctctttcttcctctagggtgtcgttaattacccgtactaaaggtttggaaaagaaaaaagagaccgcctcgtttctttttcttcgtcgaaaaaggcaataaaaatttttatcacgtttctttttcttgaaaatttttttttttgatttttttctctttcgatgacctcccattgatatttaagttaataaacggtcttcaatttctcaagtttcagtttcaaaaatctgaacccttgttcatttatgaatcatgattcatttttcttgttctattacaactttttttacttcttgctcattagaaagaaagcatagcaatctaatctaagttttaattacaaa | *S. cerevisiae* genomic DNA |
| *CCW12-pcaVo* | cacccatgaaccacacggttagtccaaaaggggcagttcagattccagatgcgggaattagcttgctgccaccctcacctcactaacgctgcggtgtgcggatacttcatgctatttatagacgcgcgtgtcggaatcagcacgcgcaagaaccaaatgggaaaatcggaatgggtccagaactgctttgagtgctggctattggcgtctgatttccgttttgggaatcctttgccgcgcgcccctctcaaaactccgcacaagtcccagaaagcgggaaagaaataaaacgccaccaaaaaaaaaaaaataaaagccaatcctcgaagcgtgggtggtaggccctggattatcccgtacaagtatttctcaggagtaaaaaaaccgtttgttttggaatttcccatttcgcggccacctacgccgctatctttgcaacaactatctgcgataactcagcaaattttgcatattcgtgttgcagtattgcgataatgggagtcttacttccaacataacggcagaaagaaatgtgagaaaattttgcatcctttgcctccgttcaagtatataaagtcggcatgctatactcagtgccctgactatctacattgttctaattattcttattctcctttattctttcctaacataccaagaaattaatcttctgtcattcgcttaaacactatatcaataa | *S. cerevisiae* genomic DNA |
| *PGK1-pcaVo* | acgcacagatattataacatctgcataataggcatttgcaagaattactcgtgagtaaggaaagagtgaggaactatcgcatacctgcatttaaagatgccgatttgggcgcgaatcctttattttggcttcaccctcatactattatcagggccagaaaaaggaagtgtttccctccttcttgaattgatgttaccctcataaagcacgtggcctcttatcgagaaagaaattaccgtcgctcgtgatttgtttgcaaaaagaacaaaactgaaaaaacccagacacgctcgacttcctgtcttcctattgattgcagcttccaatttcgtcacacaacaaggtcctagcgacggctcacaggttttgtaacaagcaatcgaaggttctggaatggcgggaaagggtttagtaccacatgctatgatgcccactgtgatctccagagcaaagttcgttcgatcgtactgttactctctctctttcaaacagaattgtccgaatcgtgtgacaacaacagcctgttctcacacactcttttcttctaaccaagggggtggtttagtttagtagaacctcgtgaaacttacatttacatatatataaacttgcataaatatactcagtgccctgactattatttggtcttttctaattcgtagtttttcaagttcttagatgctttctttttctcttttttacagatcatcaaggaagtaattatctactttttacaacaaatataaaaca | *S. cerevisiae* genomic DNA |
| *TEF1-pcaVo* | *cttcatcggtatcttcgctatattctttttagtcgaatttgcggggagaagatggatctatgctaaactaaataggcatttgaaaaacgacgacgagttacacgacatatcgccatctttaaatgagcaaccacactgggacctcatagaggacgggtctcgctggagtaaatttttcaacgggataattaagacgacaagaaggttcacgaaatctttaatgaggtctttagtcagaggcaggaacagccgtcaagggggcataagactacggtcatccccatctgcctcttcgtccagccttgccaacagggagttcttcagagacatggaggctcaaaacgaaattattgacagcctagacatcaatagtcatacaacagaaagcgaccacccaactttggctgataatagcgtataaacaatgcatactttgtacgttcaaaatacaatgcagtagatatatttatgcatattacatataatacatatcacataggaagcaacaggcgcgttggacttttaattttcgaggaccgcgaatccttacatcacacccaatcccccacaagtgatcccccacacaccatagcttcaaaatgtttctactccttttttactcttccagattttctcggactccgcgcatcgccgtaccacttcaaaacacccaagcacagcatactaaatttcccctctttcttcctctagggtgtcgttaattacccgtactaaaggtttggaaaagaaaaaagagaccgcctcgtttctttttcttcgtcgaaaaaggcaataaaaatttttatcacgtttctttttcttgaaaatttttttttttgatttttttctctttcgatgacctcccattgatatttaagttaataaacggtcttcaatttctcaagtttcagtttcaatactcagtgccctgactattttttcttgttctattacaactttttttacttcttgctcattagaaagaaagcatagcaatctaatctaagttttaattacaaa* | *S. cerevisiae* genomic DNA |
| *CCW12-pcavOO* | *cacccatgaaccacacggttagtccaaaaggggcagttcagattccagatgcgggaattagcttgctgccaccctcacctcactaacgctgcggtgtgcggatacttcatgctatttatagacgcgcgtgtcggaatcagcacgcgcaagaaccaaatgggaaaatcggaatgggtccagaactgctttgagtgctggctattggcgtctgatttccgttttgggaatcctttgccgcgcgcccctctcaaaactccgcacaagtcccagaaagcgggaaagaaataaaacgccaccaaaaaaaaaaaaataaaagccaatcctcgaagcgtgggtggtaggccctggattatcccgtacaagtatttctcaggagtaaaaaaaccgtttgttttggaatttcccatttcgcggccacctacgccgctatctttgcaacaactatctgcgataactcagcaaattttgcatattcgtgttgcagtattgcgataatgggagtcttacttccaacataacggcagaaagaaatgtgagaaaattttgcatcctttgcctccgttcaagtatataaagtcggcatgctatactcagtgccctgactatatactcagtgccctgactatttattctcctttattctttcctaacataccaagaaattaatcttctgtcattcgcttaaacactatatcaataa* | *S. cerevisiae* genomic DNA |
| *PGK1-pcavOO* | acgcacagatattataacatctgcataataggcatttgcaagaattactcgtgagtaaggaaagagtgaggaactatcgcatacctgcatttaaagatgccgatttgggcgcgaatcctttattttggcttcaccctcatactattatcagggccagaaaaaggaagtgtttccctccttcttgaattgatgttaccctcataaagcacgtggcctcttatcgagaaagaaattaccgtcgctcgtgatttgtttgcaaaaagaacaaaactgaaaaaacccagacacgctcgacttcctgtcttcctattgattgcagcttccaatttcgtcacacaacaaggtcctagcgacggctcacaggttttgtaacaagcaatcgaaggttctggaatggcgggaaagggtttagtaccacatgctatgatgcccactgtgatctccagagcaaagttcgttcgatcgtactgttactctctctctttcaaacagaattgtccgaatcgtgtgacaacaacagcctgttctcacacactcttttcttctaaccaagggggtggtttagtttagtagaacctcgtgaaacttacatttacatatatataaacttgcataaatatactcagtgccctgactatatactcagtgccctgactatgtagtttttcaagttcttagatgctttctttttctcttttttacagatcatcaaggaagtaattatctactttttacaacaaatataaaaca | *S. cerevisiae* genomic DNA |
| *TEF1-pcavOO* | cttcatcggtatcttcgctatattctttttagtcgaatttgcggggagaagatggatctatgctaaactaaataggcatttgaaaaacgacgacgagttacacgacatatcgccatctttaaatgagcaaccacactgggacctcatagaggacgggtctcgctggagtaaatttttcaacgggataattaagacgacaagaaggttcacgaaatctttaatgaggtctttagtcagaggcaggaacagccgtcaagggggcataagactacggtcatccccatctgcctcttcgtccagccttgccaacagggagttcttcagagacatggaggctcaaaacgaaattattgacagcctagacatcaatagtcatacaacagaaagcgaccacccaactttggctgataatagcgtataaacaatgcatactttgtacgttcaaaatacaatgcagtagatatatttatgcatattacatataatacatatcacataggaagcaacaggcgcgttggacttttaattttcgaggaccgcgaatccttacatcacacccaatcccccacaagtgatcccccacacaccatagcttcaaaatgtttctactccttttttactcttccagattttctcggactccgcgcatcgccgtaccacttcaaaacacccaagcacagcatactaaatttcccctctttcttcctctagggtgtcgttaattacccgtactaaaggtttggaaaagaaaaaagagaccgcctcgtttctttttcttcgtcgaaaaaggcaataaaaatttttatcacgtttctttttcttgaaaatttttttttttgatttttttctctttcgatgacctcccattgatatttaagttaataaacggtcttcaatttctcaagtttcagtttcaatactcagtgccctgactatatactcagtgccctgactattttttcttgttctattacaactttttttacttcttgctcattagaaagaaagcatagcaatctaatctaagttttaattacaaa | *S. cerevisiae* genomic DNA |
| *CYC1trunc-benO* | tcgagcagatccgccaggcgtgtatatatagcgtggatggccaggcaactttagtgctgacacataatactccataggtattttattatacaaataatgtgtttgaacttattaaaacattcttttaaggtataaacaacaggcatatatatatgtgtgcgacgacacatgatcatatggcatgcatgtgctctgtatgtatataaaactcttgttttcttcttttctctaaatattctttccttatacattaggacctttgcagcataaattactatacttctatagacacacaaacacaaatacacacactaaattaataatg | *S. cerevisiae* genomic DNA |
| **Terminator ID** | **sequence** |  |
| *ENO2* terminator | agtgcttttaactaagaattattagtcttttctgcttattttttcatcatagtttagaacactttatattaacgaatagtttatgaatctatttaggtttaaaaattgatacagttttataagttactttttcaaagactcgtgctgtctattgcataatgcactggaaggggaaaaaaaaggtgcacacgcgtggctttttcttgaatttgcagtttgaaaaataactacatggatgataagaaaacatggagtacagtcactttgagaaccttcaatcagctggtaacgtcttcgttaattggatactcaaaaaagatggatagcatgaatcacaagatggaaggaaatgcgggccacgaccacagtgatatgcatatgggagatggagatgatacct | *S. cerevisiae* genomic DNA |
| *ADH1* terminator | tggacttcttcgccagaggtttggtcaagtctccaatcaaggttgtcggcttgtctaccttgccagaaatttacgaaaagatggaaaagggtcaaatcgttggtagatacgttgttgacacttctaaataagcgaatttcttatgatttatgatttttattattaaataagttataaaaaaaataagtgtatacaaattttaaagtgactcttaggttttaaaacgaaaattcttattcttgagtaactctttcctgtaggtcaggttgctttctcaggtatagcatgaggtcgctcttattgaccacacctctaccggcatgc | *S. cerevisiae* genomic DNA |
| **Integration site homology arms** | **sequence** |  |
| 14 up homology | tcaatcaaagcaacccacaaatcctaggctgaatcatgatatcgatggaagcaatcaacaattttatcaagaccgcaccaaagcacgactatctgacaggcggagttcatcattctggtaatgtagacgtgttacaattaagcggcaataaagaagatggtactttagtatggagccatacttttgttgatgtagacaacaatgtgatagctaaatttgaagacgctctcgaaaaacttgaaagtttgcaccggcgctcatcctcatccacaggcaatgaagaacacgctaacgtttaaccgagaggagacacttcataatgatgtgagaaataagtgaatattgtaataattgttgggactccaatgtcaacaaaagctataatgtaggtattcagaatatactagaggttctcctcgaggatcttggaatccacaaaagggagtcgataaatctatataataaaaattactttatcttctttcgttttatacgttgtcgtttattatcctattacgttatcattcttcgcatttcagctttcattagatttgatgactgtttctcaaactttatgtcattttcttacacc | *S. cerevisiae* genomic DNA |
| 14 down homology | gaaaatgcaaccgataaaacattataaatcttcgcggttatctggcattgttattaaccaaaaaaatgccggcctattacaagctactgttcaataaatattgttgtaatgaagacggtccaactgtacaaatacagcaaactgtcatatataaggtgtcttatgtgacagcacttgcgttattgtcagccggagtatgtctttgtcgcattctgggctttttactttctgctcagaaggaagtacgaacaagaaaaaaaaatcaccaatgcttcccttttcagtattagtttcatatttgtttacgttcaaactcgtcgtttgcgcgataacctctaaaaaagtcagttacgtaactatatcaatcagagaatgcaaaaagcactatcataaaaatgtctctaggggatgtgagacatgtcaattataagaagtgatggtgtcatagtatatatatcataaatgattatcaaagtttcaatcctttgtattttctagtttagcgccaacttttgacaaaacctaaactttagataatcatcattcttacaatttttatctggatggcaataatctcctatataaagcccagataaactgtaaaaagaatccatcactatttgaaaaaaagtcatctggcacgtttaattatcagagcagaaatgatgaagggtgttagcgccgtccattgatgcgcctggtagtcatgatttacgtataactaacacatcatgaggacggc | *S. cerevisiae* genomic DNA |
| 16 up homology | ttcgtgaaacacgtgggataccggttaggtaatattacagtttccttacacacttactaaatacgcaacagagcatgttattaatcgtatatttaagtaataataacgtattgaccattcctaagactgaagcagtaccaagaaatggcactttttgtttatatgacatagtaaaagtgttcatttggcagaaaacaagggtatctggatactgcaatcctcattagttgcggagagcttttttaaagttttattacttcgataatgctttcctccagaactgggcgcgcactgaagccataaactgagtggttgtccctgtgtatattctacaatcgcctatttgaaacttctgggtgataatatcgatttagcgattgggtttctcccttctgagattctggataaatgagatgctaccctcacagtaaaaacagtacaaaacctttgtatagttctagtcaaatatgaactccttcattcatggattgtatactattgttcttgttgaaagattcctcttgtgattttcatggtgattttaagccttttttattgaaagaattaaatattcactaggctgcgatacgatagacaaacgaagtgattgaaacccgaattaacgga | *S. cerevisiae* genomic DNA |
| 16 down homology | tgcctacgcaacactttagctgctttttttaagcagcggcagatgaggatttaaataacgcataaccaagaatatacctaatgaagtgaaggtatacaggctgttctatggacttgaataatactggctaccctggcgtcctaatgtcgttaagaaagtaacttgatatcagattaatcacaaacttggtccacaaaataccacttatctcatataatctaaatctgactaaagggtgtaggcacggaacttcggacctagaggtattgctcaatatgtatatttctcaatggttcatctaggtaagcaaggccttgaataatatacttaatcttgtctgtttcgcaacaattgctgagtacaataatgttggaataaaaatcaactatcatctactaactagtatttacgttactagtatattatcatatacggtgttagaagatgacgcaaatgatgagaaatagtcatctaaattagtggaagctgaaacgcaaggattgataatgtaataggatcaatgaatattaacatataaaatgatgataataatatttatagaattgtgtagaattgcagattcccttttatggattcctaaatcctcgaggagaacttctagtatatctacatacctaatattatagccttaatcacaatggaatcccaacaa | *S. cerevisiae* genomic DNA |
| 18 up homology | tgtgcacaaaggccataatattatgtctacagaatatactagatgtcctccctataggatatagtaatcctctaaatggaaccgatatttctacataataatattacgattattcctccttccgttttatatgtttcattatcctagcacactatcaatctttgcatttcagcttccattagatttgatgactatttctcaatctttatgttatctccttacgccgcatgtgataatatactgctagtatgactactagttgatagaagatagttgatttttactccaacaaaagtaacaatattatttagaactatagattccattttgtgcattcccatattctcgaggaaaacttttagtatattctgtagacataatattatcgcctttgtgaacaatagaatcccaacaattgtcgcaaatttaccaattttctagattgcagtcaccttttcaattaatcactagtgtttcacttgtaacattgtcgttgttgtttaacgtattctgtcccgtgccaactatgacaaaaatgcaatgatttcagcggttaaatacgaagcgcaacaagagttagcgaaaaataagtaccaccattctacgctaccattacttactgaaattagagacaactgttatctattggcagatgttcatacggggctttcaaatattgatgaaattatgtgatgtttagaagaagattcgaactgttttcagtagatttggtaactgtgcaaccataactcatgcc | *S. cerevisiae* genomic DNA |
| 18 down homology | aatcgtccccaacaaaagtgggctctcaaaattcatcacatttaaatgcatataggaagagcaacagttggtttgcatctgatgttccttaaagatttcgacataatgtgcgaagtagataaaatgggtcatttattaatagttatttcattattaaccagttgtggtacaaatgcaactaaagaaaaaaactactaaactatccgggaaatgcgccttagattgcacttcttaattcttattttcgatttttatttttcctttgataatcataaagagaaacgacgatcatttctaaagccatttctgctagtataccgttaaataagaaaaataaagccaaatattataatttttctaatgtgaatccataaatatcaaagcatgcaaaaagggaaagaagtaatgtcttggatttatatagcgtatttgtctaaggggagccagcttt | *S. cerevisiae* genomic DNA |
| 20 up homology | gccaggcgcctttatatcatataattaagacacaaaaggataaaacaaaggtgttaactattctgcatactcactatcgtaaactgtcctgcaaatcgtgtaaatatgtatttcattttttttgcagtgaaaaaaggcatgtaaaataccgcatcaagtaactctactccgcctgtggtttcaagactaacggcttgagacaaaatgggaagaaatgattgcagaaaagccatatgtgtaatagcaaaaagctggatagtgcttaccagatgtttaccttaatttcttggtgaattagagaagtacagaagttttactattaatcccaccatagaaatttgtataggaaagtagtttattggagttattggatatactgtgtaaactatttcttgaaattgtaatcttaagatgctcttcttattctattaaaaatagaaaatgattttcatatttatttatttatttatattttggcattactcttcatcatttttttccctctaagaagcttcctttctttttataaggataacaaaaccaaaaggaatattgggtcagatgaatggacgcgaatgcaagacagaagtccaaatcacgtcaagacaaagaaagaaagaaagaaaaactaacacattaatgtagttttaaaatttcaaatccgaacaacagagcatagggtttcgcaaa | *S. cerevisiae* genomic DNA |
| 20 down homology | aatggaaggtcgggatgagcatatacaagcactaagaagaacaatacagaactctacacggtattattgtgctacaagctcgagtaaaaccgagtgttttgacgatactaacgttgttaagaaagtaacttgttatcaaactcattaccaacttgtgattaattggtgaataatatgataattgtcgaaattccattgttggtaaagcctataatattatgtatacagattatactagaaattctctcgagaatataagaatccccaaaattgaatcggtatttctacatactaatattaccattacttctcctttcgttttatatgtttcattcctattacattatcgatctttgcatttcagcttccattatatttgatgtctgttttatgtccccacgttacaccgcatgtgacagtatactagtaacatgagtgctaccgaatagatgacattttagactttcattccaacaacttggttgacagaatgttacgtaccctatatctaatctatatgaggcctgaatctaactgaaaggtggaatttcagtaatttatcaagctttaataagtttgggtagtttaactgtgcaaaaaggtatttaccttacatactgaatcttgtctgtttggtagcggctgctttat | *S. cerevisiae* genomic DNA |

**Table S4. Oligonucleotides used in this work**

| **Purpose** | **Oligo ID** | **Nucleotide sequence (5’ – 3’)** | **Template** |
| --- | --- | --- | --- |
| 1132 with mut rolr overhang - tdh3 rv | MN987 | gcgctgtactagcatctttgtgttgggagggtgtaggcattcgaaactaagttcttggtgttttaaaac | *S. cerevisiae* genomic DNA |
| LTP1-TEF1fw | MN785 | ccggtcttagaaaacgcataaacatacaagtggacagatgccttgccaacagggagttct | *S. cerevisiae* genomic DNA |
| envy-TEF1 rv | MN786 | taggaactacacctgtaaacaattcctcgcctttagacattgtttttttgtaattaaaacttagattagattgctatgctt | *S. cerevisiae* genomic DNA |
| rolo-tef1 rv | MN787 | tgaatcatgattcataaatgaacaagggttcagattttcgtgataaaaatttttattgcctttttcg | *S. cerevisiae* genomic DNA |
| rolo-tef1 fwd | MN788 | aaaatctgaacccttgttcatttatgaatcatgattcatctttcgatgacctcccattga | *S. cerevisiae* genomic DNA |
| LTP1-TDH3fw | MN789 | ccggtcttagaaaacgcataaacatacaagtggacagatgtcgagtttatcattatcaatactgccatt | *S. cerevisiae* genomic DNA |
| envy-TDH3rv | MN790 | taggaactacacctgtaaacaattcctcgcctttagacattgtttttcgaaactaagttcttggtgttttaaaac | *S. cerevisiae* genomic DNA |
| rolo-tdh3 rv | MN791 | tgaatcatgattcataaatgaacaagggttcagatttttacctaccgtctttatatacttattagtcaag | *S. cerevisiae* genomic DNA |
| rolo-tdh3 fwd | MN792 | aaaatctgaacccttgttcatttatgaatcatgattcaaaattctacttttatagttagtcttttttttagttttaaaac | *S. cerevisiae* genomic DNA |
| PcaV fwd - tdh3 overhang | MN796 | ttttagttttaaaacaccaagaacttagtttcgaaaaacaatggcagcagtagaccttgcca | PcaV gBlock |
| PcaV rv- eno2 overhang | MN797 | aataagcagaaaagactaataattcttagttaaaagcacttcaaccaggagccacggct | PcaV gBlock |
| pcavo rv - ccw | MN800 | gaataattagaacaatgtagatagtcagggcactgagtatagcatgccgactttatatacttgaacgga | *S. cerevisiae* genomic DNA |
| pcavo fw - ccw | MN801 | gtatataaagtcggcatgctatactcagtgccctgactatctacattgttctaattattcttattctcctttattctttcctaacatac | *S. cerevisiae* genomic DNA |
| pcavo F pgk (to be used with 514) | MN823 | atactcagtgccctgactatttcgtagtttttcaagttcttagat | *S. cerevisiae* genomic DNA |
| pcavo R pgk (to be used with 526) | MN824 | atagtcagggcactgagtattttatgcaagtttatatatatgtaaatgtaag | *S. cerevisiae* genomic DNA |
| pcavo-tdh3 fwd | MN833 | atactcagtgccctgactataaattctacttttatagttagtcttttttttagttttaaaac | *S. cerevisiae* genomic DNA |
| pcavo-tdh3 rv | MN834 | atagtcagggcactgagtattacctaccgtctttatatacttattagtcaag | *S. cerevisiae* genomic DNA |
| yeastract TEF1 fwd with ltp1 overhang | MN864 | ccggtcttagaaaacgcataaacatacaagtggacagatgcttcatcggtatcttcgctatattcttttt | *S. cerevisiae* genomic DNA |
| yeastract TEF1 rv with envy overhang | MN865 | taggaactacacctgtaaacaattcctcgcctttagacattgtttttttgtaattaaaacttagattagattgctatgctttc | *S. cerevisiae* genomic DNA |
| TEF1-rolo fwd | MN866 | aaaatctgaacccttgttcatttatgaatcatgattcatttttcttgttctattacaactttttttacttcttg | *S. cerevisiae* genomic DNA |
| TEF1-rolo rv | MN867 | tgaatcatgattcataaatgaacaagggttcagatttttgaaactgaaacttgagaaattgaagacc | *S. cerevisiae* genomic DNA |
| TEF1-pcavo fwd | MN874 | tttctcaagtttcagtttcaatactcagtgccctgactattttttcttgttctattacaactttttttacttcttg | *S. cerevisiae* genomic DNA |
| TEF1-pcavo rv | MN875 | atagtcagggcactgagtattgaaactgaaacttgagaaattgaagacc | *S. cerevisiae* genomic DNA |
| gRNA FgF20 fw hyb | MN1034 | tcctcgctggcgccggctgggcaacaccttcgggtggcgaatgggactttgttagagctgttacaagtta | n/a |
| gRNA FgF20 rv hyb | MN1035 | gttgataacggactagccttattttaacttgctatttctagctctaaaactaacttgtaacagctctaac | n/a |
| Fgf20 up reverse LV3 overhang use with GC1764 | MN1036 | gcatttttattatataagttgttttattcagagtattccttttgcgaaaccctatgctct | *S. cerevisiae* genomic DNA |
| TEF1 fwd lv3 overhang | MN1037 | aggaatactctgaataaaacaacttatataataaaaatgccttcatcggtatcttcgctatattct | *S. cerevisiae* genomic DNA |
| FgF20 down fwd use with GC1767 | MN1038 | cctctttatattacatcaaaataagaaaataattataacaaatggaaggtcgggatgag | *S. cerevisiae* genomic DNA |
| Adh1t fwd with bleoR overhang use with payge124 | MN1039 | tgttcattttgttgctgaagaacaagattaatggacttcttcgccag | *S. cerevisiae* genomic DNA |
| bleoR Rv with ADH1t overhang | MN1040 | gattggagacttgaccaaacctctggcgaagaagtccattaatcttgttcttcagcaaca | pYTK080 |
| tef1 lv3 overhang | MN1042 | aggaatactctgaataaaacaacttatataataaaaatgccttcatcggtatcttcgctatattct | *S. cerevisiae* genomic DNA |
| tef1 rv blunt | MN1043 | tgtttttttgtaattaaaacttagattagattgctatgct | *S. cerevisiae* genomic DNA |
| bleoR forward add overhang pgk | MN1047 | gaagtaattatctactttttacaacaaatataaaacaatggctaaattaacatctgccgt | pYTK080 |
| bleoR forward add overhang ccw | MN1048 | ctgtcattcgcttaaacactatatcaataaaaaacaatggctaaattaacatctgccgt | pYTK080 |
| bleoR fwd blunt | MN1060 | atggctaaattaacatctgccgt | pYTK080 |
| 18 grna hyb fwd | MN1063 | tcctcgctggcgccggctgggcaacaccttcgggtggcgaatgggactttatagaattactattgaagag | n/a |
| 18 grna hyb rv | MN1064 | gttgataacggactagccttattttaacttgctatttctagctctaaaacctcttcaatagtaattctat | n/a |
| pcavo2x Tef1OF | MN1136 | atactcagtgccctgactatatactcagtgccctgactattttttcttgttctattacaactttttttacttcttg | *S. cerevisiae* genomic DNA |
| pcavo2x Tef1OR | MN1137 | atagtcagggcactgagtatatagtcagggcactgagtattgaaactgaaacttgagaaattgaagacc | *S. cerevisiae* genomic DNA |
| pcavo2x TDH3 OF | MN1138 | atactcagtgccctgactatatactcagtgccctgactataaattctacttttatagttagtcttttttttagttttaaaac | *S. cerevisiae* genomic DNA |
| pcavo2x TDH3 OR | MN1139 | atagtcagggcactgagtatatagtcagggcactgagtattacctaccgtctttatatacttattagtcaag | *S. cerevisiae* genomic DNA |
| bleoR F cyc1 OH | MN1140 | atagacacacaaacacaaatacacacactaaattaataatgaaaacaatggctaaattaacatctgccgt | pYTK080 |
| 18up F n | MN1143 | tgtgcacaaaggccataatattatgtctacagaat | *S. cerevisiae* genomic DNA |
| 18down R n | MN1144 | aaagctggctccccttagacaaatacgcta | *S. cerevisiae* genomic DNA |
| RolR F tdh3 overhang | MN1167 | ttttagttttaaaacaccaagaacttagtttcgaaaaacaatgcctacaccctcccagca | *S. cerevisiae* genomic DNA |
| RolR R eno overhang | MN1168 | aataagcagaaaagactaataattcttagttaaaagcactctactgcggcttaaccgggc | *S. cerevisiae* genomic DNA |
| gap promoter f aid6 | MN1236 | cattatcaatactgccatttcaaaga | pAID6 |
| lbcas rv - 1.1 kb product aid6 | MN1237 | attctccttctcggttctgg | pAID6 |
| tef1promf | MN1238 | atagcttcaaaatgtttctactccttt | pAID6 |
| sacas rv - 1.1 kb product | MN1239 | attccttgatgtccttccac | pAID6 |
| ura3 grna f coding sequence 1 | MN1244 | tcctcgctggcgccggctgggcaacaccttcgggtggcgaatgggacttttatcttgactgatttttcca | n/a |
| ura3 grna r coding sequence 1 | MN1245 | gttgataacggactagccttattttaacttgctatttctagctctaaaactggaaaaatcagtcaagata | n/a |
| ura3 grna f coding sequence 2 | MN1246 | tcctcgctggcgccggctgggcaacaccttcgggtggcgaatgggacttttaactccagtaattccttgg | n/a |
| ura3 grna r coding sequence 2 | MN1247 | gttgataacggactagccttattttaacttgctatttctagctctaaaacccaaggaattactggagtta | n/a |
| eno2rv bunt | MN1248 | aggtatcatctccatctccc | *S. cerevisiae* genomic DNA |
| ura3 down f with eno2 overhang use with 1195 | MN1249 | gaccacagtgatatgcatatgggagatggagatgatacctaaaactgtattataagtaaatgcatgtatac | *S. cerevisiae* genomic DNA |
| Pcavo 2x ccw F | MN1120 | atactcagtgccctgactatatactcagtgccctgactatttattctcctttattctttcctaaca | *S. cerevisiae* genomic DNA |
| Pcavo 2x ccw R | MN1121 | atagtcagggcactgagtatatagtcagggcactgagtatagcatgccgactttatatactt | *S. cerevisiae* genomic DNA |
| Pcavo 2x pgk F | MN1122 | atactcagtgccctgactatatactcagtgccctgactatgtagtttttcaagttcttagatgcttt | *S. cerevisiae* genomic DNA |
| Pcavo 2x pgk R | MN1123 | atagtcagggcactgagtatatagtcagggcactgagtatatttatgcaagtttatatatatgtaaatgtaagtt | *S. cerevisiae* genomic DNA |
| PaAroZ F TDH3 overhang | MN960 | ttttagttttaaaacaccaagaacttagtttcgaaaaacaatgccatccaagttggccat | DT64 genomic DNA |
| PaAroZ R ADH1 overhang | MN961 | ttgattggagacttgaccaaacctctggcgaagaagtccattacaaggcagcggataaagaca | DT64 genomic DNA |
| PaAroZ F PGK1 overhang | MN962 | aaggaagtaattatctactttttacaacaaatataaaacaatgccatccaagttggccat | DT64 genomic DNA |
| ADH1t fwd with aroz overhang | MN963 | atcagaatgttgtctttatccgctgccttgtaatggacttcttcgccaga | *S. cerevisiae* genomic DNA |
| envy f blunt | MN1424 | atgtctaaaggcgaggaattgttta | *S. cerevisiae* genomic DNA |
| PAD1 native promoter gRNA fwd | MN1425 | tcctcgctggcgccggctgggcaacaccttcgggtggcgaatgggactttattgaagctgctcaatgcgt | n/a |
| PAD1 native promoter gRNA rv | MN1426 | gttgataacggactagccttattttaacttgctatttctagctctaaaacacgcattgagcagcttcaat | n/a |
| pad1 ko donor 1 fwd | MN1427 | tcattagtagcacttcgtatcg | *S. cerevisiae* genomic DNA |
| pad1 ko donor 1 rv | MN1428 | actccgtagattgaaatcaatctat | *S. cerevisiae* genomic DNA |
| pad1 donor 2 fwd | MN1429 | acataatgctgcaaatatagattgatttcaatctacggagtcactttttctgagcattttattacgtt | *S. cerevisiae* genomic DNA |
| pad1 donor 2 rv | MN1430 | taagctgacattgaaaatgagtaaga | *S. cerevisiae* genomic DNA |
| aro2 grna fwd | MN1431 | tcctcgctggcgccggctgggcaacaccttcgggtggcgaatgggactttagtcttgccgaactcggtac | n/a |
| aro2 grna rv | MN1432 | gttgataacggactagccttattttaacttgctatttctagctctaaaacgtaccgagttcggcaagact | n/a |
| aro2 up fwd | MN1433 | atacagaccattcagcttcag | *S. cerevisiae* genomic DNA |
| aro2 up rv | MN1434 | gcctctgtgctatgatactattt | *S. cerevisiae* genomic DNA |
| aro2 down fwd | MN1435 | gtatagtatcgaaaaaagaaaatagtatcatagcacagaggcagatagaaacaaaacccctcaa | *S. cerevisiae* genomic DNA |
| aro2 down rv | MN1436 | gtcgtgtcagaaacaaggatat | *S. cerevisiae* genomic DNA |
| aro2 cyc1 donor f | MN1448 | gtatagtatcgaaaaaagaaaatagtatcatagcacagatgctcgagcagatccgcca | *S. cerevisiae* genomic DNA |
| aro2 cyc1 donor r | MN1449 | atatgtggtgacgcggaacagtttcccaaacgttgacattgttttcattattaatttagtgtgtgtatttgtgt | *S. cerevisiae* genomic DNA |
| CYC1a guide Fw | MD527 | cagtgaggtctcatgatcaatttctactaagtgtag | n/a |
| CYC1a guide Rv | MD528 | gtcgtaggtctcaacttgcgcttgatccaccaaccaacgctcgcatctacacttagtagaaattgatcatg | n/a |
| tRNA_GLY_ site 1-2 Fw | MD529 | cagtgaggtctcaaagtggtttagtggtaaaatccaacgttgccatc | n/a |
| tRNA_GLY_ site 1-2 Rv | MD530 | gtcgtaggtctcatgcgcaagcccggaatcgaaccgggggcccaacgatggcaacgttggattttac | n/a |
| tRNA_GLY_ site 2-3 Fw | MD533 | cagtgaggtctcagcaagtggtttagtggtaaaatccaacgttgccatc | n/a |
| tRNA_GLY_ site 2-3 Rv | MD534 | gtcgtaggtctcacaagcccggaatcgaaccgggggcccaacgatggcaacgttggattttac | n/a |
| pGG1 backbone amplification Fw | AK1006 | cggctaaggtctcatgaagttttagtactctgtaattttaggtatgag | *pGG1* |
| pGG1 backbone amplification Rv | AK1007 | ggctacggtctctatccaccaaccaacgctcg | *pGG1* |
| Part 1 pAK091 amplification Fw | AK1008 | ggctaaggtctcaggatcaagcgcaagtggtttagtggtaaaatccaacgttgccatc | n/a |
| Part 1 pAK091 amplification Rv | AK1009 | ggctacggtctctaaacttgcctccgttcaagtatattgcgcaagcccggaatcgaaccgggggcccaacgatggcaacgttggattttaccac | n/a |
| Part 2 pAK091 amplification Fw | AK1010 | taaggtctcagttttagagctagaaatagcaagttaaaataaggctagtccgttatcaacttgaaaaagtggcaccgagtcggtgcttttttgatccgcg | n/a |
| Part 2 pAK091 amplification Rv | AK1011 | tacggtctcttgtgtgcgcaagcccggaatcgaaccgggggcccaacgatggcaacgttggattttaccactaaaccacttgcgcggatcaaaaaagcac | n/a |
| Part 3 pAK091 amplification Fw | AK1012 | ggctaaggtctcacacaccaaatataccacaaccgggaaaagatttgattgcattttctgccacggaactctagcaggcgcataacataagtca | n/a |
| Part 3 pAK091 amplification Rv | AK1013 | ggctacggtctctttcaacttaaggcgaagttgttccacaaggacaatatttgtgacttatgttatgcgcctgc | n/a |
| CYC1a amplification Fw | AK1062 | ggctaaggtctcatgataatttctactaagtgtagatgcgagcgttggttggtggatcaagttttgagaccttagcc | n/a |
| CYC1a amplification Rv | AK1063 | ggctaaggtctcaaaacttgatccaccaaccaacgctcgcatctacacttagtagaaattatcatgagaccttagcc | n/a |
| CCW12i amplification Fw | AK1064 | ggctaaggtctcatgatatatacttgaacggaggcaagttttagagctagaaatagcaagttaaaataaggctagtccgttatc | n/a |
| CCW12i amplification Rv | AK1065 | ggctaaggtctcaaaacggatcaaaaaagcaccgactcggtgccactttttcaagttgataacggactagccttattttaac | n/a |
| ADE2d amplification Fw | AK1066 | ggctaaggtctcatgatcacaccaaatataccacaaccg | *pGG2* |
| ADE2d amplification Rv | AK1067 | ggctaaggtctcaaaacggatcaaaaaaatctcgccaac | *pGG2* |
| PCR diag USER-XII-1 Fw | AK727 | cttcttggacttcttaccacc | *S. cerevisiae* genomic DNA |
| PCR diag USER-XII-1 Rv | AK728 | cagtttcatccaatacgaacg | *S. cerevisiae* genomic DNA |
| P_CYC1_ amplification Fw | MD537 | aggaatactctgaataaaacaacttatataataaaaatgcatttttttggaaaaccaag | *S. cerevisiae* genomic DNA |
| P_CYC1_ amplification Rv | MD540 | cagcttcacctttactaaccatcattattaatttagtgtgtgtatttg | *S. cerevisiae* genomic DNA |
| *mScarlet* amplification Fw | MD542 | atacacacactaaattaataatgatggttagtaaaggtgaagctg | *pBBK35 (Bean et al., 2022)* |
| *mScarlet* amplification Rv | MD538 | tgttataattattttcttattttgatgtaatataaagaggataaaattaaagtagcagtacttcaac | *pBBK35 (Bean et al., 2022)* |
| USER XII-1 5’-Homology Fw | MD370 | ctttagacaattgttgctttggag | *S. cerevisiae* genomic DNA |
| USER XII-1 5’-Homology Rv | MD371 | gcatttttattatataagttgttttattcagagtattcctgaaagaaccgaaccgatgc | *S. cerevisiae* genomic DNA |
| USER XII-1 3’-Homology Fw | MD372 | cctctttatattacatcaaaataagaaaataattataacacttcccgtgaatcaactgc | *S. cerevisiae* genomic DNA |
| USER XII-1 3’-Homology Rv | MD373 | gtgaaatcttcaccaaatgctc | *S. cerevisiae* genomic DNA |
| pBB95 no Cas plasmid backbone Fw | AK839 | gggtggcgaatgggacttt | pBB95 (Bean et al., 2022) |
| pBB95 no Cas plasmid Backbone Rv | AK842 | agctctcgagaacccttaatgtcgacagtctgagcaaaaggccagcaaaag | pBB95 (Bean et al., 2022) |
| XII-1 SaCas9 deletion guide Fw | AK836 | atactgtcagaggtggcattttacataacaatagtgacaaaaaataaaggatcaaaaaaatctcgccaac | pAK091 |
| XII-1 SaCas9 deletion guide Rv | AK837 | ctcgctggcgccggctgggcaacaccttcgggtggcgaatgggactttgcatcgcaagtcaggggccaggttttagtactctgtaattttaggtatgag | pAK091 |
| P_CYC1o_ amplification Fw | AK1162 | tgataaattgaatttcattgcaatggtgattcgagcagatccgccaggcg | MN3037 genomic DNA |
| P_CYC1o_ amplification Rv | AK1075 | gcagatgttaatttagccattattaatttagtgtgtgtatttgtgtttg | *MN3037 genomic DNA* |
| *bleoR* amplification Fw | AK1076 | atacacacactaaattaataatggctaaattaacatctgcc | MN3034 genomic DNA |
| *bleoR* amplification Rv | AK1077 | cgatttcaattcaattcaatttaatcttgttcttcagcaacaaaatg | *MN3034 genomic DNA* |
| T_PGK1_ amplification Fw | AK1078 | ttgctgaagaacaagattaaattgaattgaattgaaatcgatagatc | *S. cerevisiae* genomic DNA |
| T_PGK1_ amplification Rv | AK1163 | gataatataccagtaatattaataccagtgacatagaaatatcgaatgggaaaaaaaaac | *S. cerevisiae* genomic DNA |
| FgF14 5’-Homology Fw | AK1150 | tatggagccatacttttgttgatgtag | *S. cerevisiae* genomic DNA |
| FgF14 5’-Homology Rv | AK1151 | atcaccattgcaatgaaattcaatttatc | *S. cerevisiae* genomic DNA |
| FgF14 3’-Homology Fw | AK1152 | cactggtattaatattactggtatattatctaatatg | *S. cerevisiae* genomic DNA |
| FgF14 3’-Homology Rv | AK1153 | ttctgagcagaaagtaaaaagcccag | *S. cerevisiae* genomic DNA |
| FgF14 SpCas9 cut guide Fw | AK1148 | gactatcattgtgattttatttgt | n/a |
| FgF14 SpCas9 cut guide Rv | AK1149 | aaacacaaataaaatcacaatgat | n/a |
| URA3 5’-Homology Fw | AK1129 | tatgttaattaccttttttgcgagg | *S. cerevisiae* genomic DNA |
| URA3 5’-Homology Rv | AK1130 | gatttatcttcgtttcctgcagg | *S. cerevisiae* genomic DNA |
| URA3 3’-Homology Fw | AK1131 | tcttaacccaactgcacagaacaaaaacctgcaggaaacgaagataaatcaaaactgtattataagtaaatgcatgtatac | *S. cerevisiae* genomic DNA |
| URA3 3’-Homology Rv | AK1132 | tgtgaaacatctttctaccaga | *S. cerevisiae* genomic DNA |
| URA3 SpCas9 cut guide Fw | AK1119 | tcctcgctggcgccggctgggcaacaccttcgggtggcgaatgggacttttatcttgactgatttttcca | n/a |
| URA3 SpCas9 cut guide Rv | AK1120 | gttgataacggactagccttattttaacttgctatttctagctctaaaactggaaaaatcagtcaagata | n/a |
| PAD1 5’-Homology Fw | AK1123 | tcattagtagcacttcgtatcg | *S. cerevisiae* genomic DNA |
| PAD1 5’-Homology Rv | AK1124 | actccgtagattgaaatcaatctat | *S. cerevisiae* genomic DNA |
| PAD1 3’-Homology Fw | AK1125 | acataatgctgcaaatatagattgatttcaatctacggagtcactttttctgagcattttattacgtt | *S. cerevisiae* genomic DNA |
| PAD1 3’-Homology Rv | AK1126 | taagctgacattgaaaatgagtaaga | *S. cerevisiae* genomic DNA |
| PAD1 SpCas9 cut guide Fw | AK1135 | tcctcgctggcgccggctgggcaacaccttcgggtggcgaatgggactttattgaagctgctcaatgcgt | n/a |
| PAD1 SpCas9 cut guide Rv | AK1136 | gttgataacggactagccttattttaacttgctatttctagctctaaaacacgcattgagcagcttcaat | n/a |
| SOE1_i.F | MT637 | ATCAGGTCTCATGATATACTAGGGTATTCTCCTCAG | p426*-SpSgH |
| RVS167_i.F | MT638 | ATCAGGTCTCATGATCGTTCTGCATCCTCGTAAAC | p426*-SpSgH |
| DUG3_i.F | MT639 | ATCAGGTCTCATGATTCATTGCTCAAATTATCCGC | p426*-SpSgH |
| RIM4_i.F | MT640 | ATCAGGTCTCATGATATTAGAGGGAGGATCACGAAG | p426*-SpSgH |
| SpCasScaff_GG.R | MT641 | ATCGGTCTCCAAACGGATCAAAAAAGCACCGAC | p426*-SpSgH |
| PDR3_a.F | MT642 | ATCGGTCTCATGATAATTTCTACTAAGTGTAGATCGCGGAGCTG | n/a |
| PDR3_a.R | MT643 | ACTGGTCTCAAAACGCAGAGGAGGCAGCTCCGCGATCTAC | n/a |
| CCS1_a.F | MT644 | ACTGGTCTCATGATAATTTCTACTAAGTGTAGATTTCTCTAATAAGCTC | n/a |
| CCS1_a.R | MT645 | ACTGGTCTCAAAACCCAACGAGCTTATTAGAGAAATCTACACTTAG | n/a |
| RNH202_a.F | MT646 | ACTGGTCTCATGATAATTTCTACTAAGTGTAGATAGTATGGCG | n/a |
| RNH202_a.R | MT647 | ACTGGTCTCAAAACCGCTACGAAGAAACGCCATACTATCTACACTTAGTAG | n/a |
| PDR3_tGly.R | MT648 | ATCAGGTCTCAGCAACGTTGGATTTTACCACTAAACCACTTGCAGAGGAGGCAGCTCCGCGATCTAC | n/a |
| tGly_RNH202.F | MT649 | ACTAGGTCTCTTGCCATCGTTGGGCCCCCGGTTCGATTCCGGGCTTGCGCAAATTTCTACTAAGTGTAGATAGTATGGCG | n/a |
| RNH202_tGly.R | MT650 | ACTGGTCTCAGCAACGTTGGATTTTACCACTAAACCACTTCGCTACGAAGAAACGCCATACTATCTACACTTAGTAG | n/a |
| tGly_DUG3.F | MT651 | ACTGGTCTCATTGCCATCGTTGGGCCCCCGGTTCGATTCCGGGCTTGCGCATCATTGCTCAAATTATCCGC | p426*-SpSgH |
| CCS1_tGly.R | MT652 | ACTGGTCTCAGCAACGTTGGATTTTACCACTAAACCACTTCCAACGAGCTTATTAGAGAAATCTACACTTAG | n/a |
| tGly_RIM4.F | MT653 | ACTGGTCTCATTGCCATCGTTGGGCCCCCGGTTCGATTCCGGGCTTGCGCAATTAGAGGGAGGATCACGAAG | p426*-SpSgH |
| tGly_CCS1.F | MT654 | ACTGGTCTCATTGCCATCGTTGGGCCCCCGGTTCGATTCCGGGCTTGCGCAAATTTCTACTAAGTGTAGATTTCTCTAATAAGCTC | n/a |
| iIDP1_M3.F | MT775 | ATCAGGTCTCATGATTCCACTTGCCCATATAAAGAG | p426*-SpSgH |
| iTP1_M3.F | MT776 | ATCAGGTCTCATGATTGTTACAGCAGCCGAG | p426*-SpSgH |
| iTP1_tGly_m3 | MT777 | ATCAGGTCTCAGCAACGTTGGATTTTACCACTAAACCACTTgcgcGGATCAAAAAAGCACCGAC | n/a |
| tGly_iTP1_M3..F | MT778 | ACTGGTCTCATTGCCATCGTTGGGCCCCCGGTTCGATTCCGGGCTTGCGCATGTTACAGCAGCCGAG | p426*-SpSgH |
| tGly_iIDP1_M3.F | MT779 | ACTGGTCTCATTGCCATCGTTGGGCCCCCGGTTCGATTCCGGGCTTGCGCATCCACTTGCCCATATAAAGAG | p426*-SpSgH |
| cyc1trunc fwd idp1 | MN.LMSE.1099 | gaaatcttccttcaagcaattgtgagacaacagacgcacatcgagcagatccgcca | *S. cerevisiae* genomic DNA |
| cyc1trunc rv idp1 | MN.LMSE.1100 | ggcgagaggtggaaaataatcttctagataacatactcatTGTTTTcattattaatttagtgtgtgtatttgtgt | *S. cerevisiae* genomic DNA |
| cyc1trunc fwd rvs167 | MN.LMSE.1103 | taaagtgaggctcacctggtcatttaacaccaagaatcatcgagcagatccgcca | *S. cerevisiae* genomic DNA |
| cyc1trunc rv rvs167 | MN.LMSE.1104 | gagccctactaacagccttggtaaaccctttaaaactcatTGTTTTcattattaatttagtgtgtgtatttgtgt | *S. cerevisiae* genomic DNA |
| cyc1trunc fwd rim4 | MN.LMSE.1107 | aagaaagaaaaaaatattatcttttttagcttgttgaaactcgagcagatccgcca | *S. cerevisiae* genomic DNA |
| cyc1trunc rv rim4 | MN.LMSE.1108 | gatcacgaagggaatccgcagtgctgatttcTGTtttcatTGTTTTcattattaatttagtgtgtgtatttgtgt | *S. cerevisiae* genomic DNA |
| cyc1trunc fwd tpi1 | MN.LMSE.1111 | ttaaatctataactacaaaaaacacatacataaactaaaatcgagcagatccgcca | *S. cerevisiae* genomic DNA |
| cyc1trunc rv tpi1 | MN.LMSE.1112 | ttcTTTaatggattgtttggaaccgtttaatttaaagttaccaccgacaaagaaagttctagccatTGTTTTcattattaatttagtgtgtgtatttgtg | *S. cerevisiae* genomic DNA |
| cyc1trunc fwd pdr3a | MN.LMSE.1115 | actgcatcagcagttttattaattttttcttattgcgtgacgagtttatcattatcaatactgc | *S. cerevisiae* genomic DNA |
| cyc1trunc rv pdr3a | MN.LMSE.1116 | ctgtcgaaacttttgatctagttgatttcttcactttcatTGTTTTtcgaaactaagttcttggtgt | *S. cerevisiae* genomic DNA |
| cyc1trunc fwd ccs1a | MN.LMSE.1119 | agactaagaaaagaataaagatatctttgcaaggcagaaacgagtttatcattatcaatactgc | *S. cerevisiae* genomic DNA |
| cyc1trunc rv ccs1a | MN.LMSE.1120 | gaatggcataagtagcctcgtatgtatcgttcgtggtcatTGTTTTtcgaaactaagttcttggtgt | *S. cerevisiae* genomic DNA |
| idp1 grna f | MN.LMSE.1097 | CGCCGGCTGGGCAACACCTTCGGGTGGCGAATGGGACTTTtgtgagacaacagacgcaca | n/a |
| Idp1 grna r | MN.LMSE.1098 | GACTAGCCTTATTTTAACTTGCTATTTCTAGCTCTAAAACtgtgcgtctgttgtctcaca | n/a |
| rvs167 grna f | MN.LMSE.1101 | CGCCGGCTGGGCAACACCTTCGGGTGGCGAATGGGACTTTtcatttaacaccaagaatca | n/a |
| rvs167 grna r | MN.LMSE.1102 | GACTAGCCTTATTTTAACTTGCTATTTCTAGCTCTAAAACtgattcttggtgttaaatga | n/a |
| rim4 grna f | MN.LMSE.1105 | CGCCGGCTGGGCAACACCTTCGGGTGGCGAATGGGACTTTggaatccgcagtgctgattt | n/a |
| rim4 grna r | MN.LMSE.1106 | GACTAGCCTTATTTTAACTTGCTATTTCTAGCTCTAAAACaaatcagcactgcggattcc | n/a |
| tpi grna f | MN.LMSE.1109 | CGCCGGCTGGGCAACACCTTCGGGTGGCGAATGGGACTTTcggttccaaacaatccatta | n/a |
| tpi grna r | MN.LMSE.1110 | GACTAGCCTTATTTTAACTTGCTATTTCTAGCTCTAAAACtaatggattgtttggaaccg | n/a |
| pdr3a grna f | MN.LMSE.1113 | CGCCGGCTGGGCAACACCTTCGGGTGGCGAATGGGACTTTtgatttcttcactttcattg | n/a |
| pdr3a grna r | MN.LMSE.1114 | GACTAGCCTTATTTTAACTTGCTATTTCTAGCTCTAAAACcaatgaaagtgaagaaatca | n/a |
| ccs1a grna f | MN.LMSE.1117 | CGCCGGCTGGGCAACACCTTCGGGTGGCGAATGGGACTTTcatttttgtgctgatcgaat | n/a |
| ccs1a grna r | MN.LMSE.1118 | GACTAGCCTTATTTTAACTTGCTATTTCTAGCTCTAAAACattcgatcagcacaaaaatg | n/a |
| cyc1 f screen | MN.LMSE.1121 | tcgagcagatccgcca | n/a |
| tdh3 f screen | MN.LMSE.1122 | cgagtttatcattatcaatactgc | n/a |
| Idp1 r screen | MN.LMSE.1123 | aagtaagatcttctttcggag | n/a |
| rvs167 r screen | MN.LMSE.1124 | ggttgtcctctggtatgg | n/a |
| rim4 r screen | MN.LMSE.1125 | gatagggctgctgctaag | n/a |
| tpi1 r screen | MN.LMSE.1126 | ctgatggtaccattaggagatt | n/a |
| pdr3 rv screen | MN.LMSE.1127 | ttgattatctagcttaatcgcag | n/a |
| ccs1 r screen | MN.LMSE.1128 | ctgaacgattcttgctagg | n/a |

**Table S5. gRNA sequences used in this work**

| Target / Purpose | Sequence |  | Source |
| --- | --- | --- | --- |
| Flagfeldt site 14 | ATCATTGTGATTTTATTTGT |  | (Flagfeldt et al., 2009) |
| Flagfeldt site 16 | CGTAGATTCCTATATCCTAC |  | Flagfeldt et al., 2009) |
| Flagfeldt site 18 | ATAGAATTACTATTGAAGAG |  | Flagfeldt et al., 2009) |
| Flagfeldt site 20 | GTTAGAGCTGTTACAAGTTA |  | Flagfeldt et al., 2009) |
| *URA3* / Deletion | TATCTTGACTGATTTTTCCA  TAACTCCAGTAATTCCTTGG |  | This work |
| *PAD1* / Deletion | ATTGAAGCTGCTCAATGCGT |  | This work |
| *ARO2* / Interference (*P_CYC1_* insertion) | AAAATAGTATCATAGCACAG |  | This work |
| *XII-1* / Deletion (*SaCas9 scaffold*) | GTGCTACTGTTGGGAAGAGCC |  | This work |
| *CYC1* / Activation (LbCas12a scaffold) | GCGAGCGTTGGTTGGTGGATCAA |  | This work |
| CCW12 / Interference (SpCas9 scaffold) | ATATACTTGAACGGAGGCAA |  | This work |
| *ADE2* / Deletion (SaCas9 scaffold) | GAACAACTTCGCCTTAAGTTGAA |  | This work |
| *IDP1* / Interference (*P_CYC1_* insertion) | tgtgagacaacagacgcaca |  | This work |
| *Rvs167* / Interference (*P_CYC1_* insertion) | tcatttaacaccaagaatca |  | This work |
| *RIM4* / Interference (*P_CYC1_* insertion) | ggaatccgcagtgctgattt |  | This work |
| *TPI1* / Interference (*P_CYC1_* insertion) | cggttccaaacaatccatta |  | This work |
| *PDR3* / Activation (*P_TDH3_* insertion) | tgatttcttcactttcattg |  | This work |
| *CCS1* / Activation (*P_TDH3_* insertion) | catttttgtgctgatcgaat |  | This work |

**Supplementary figures:**

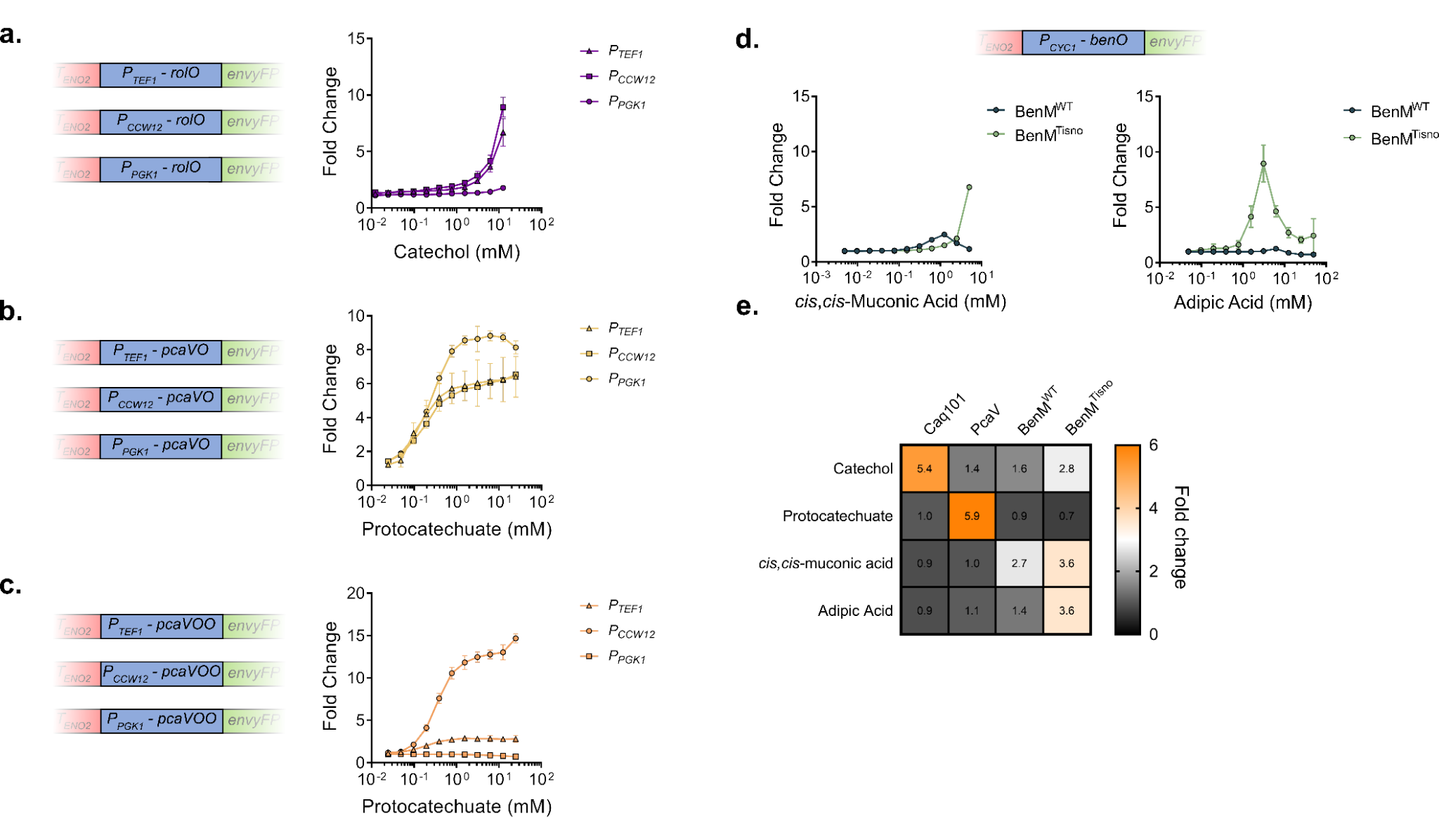

**Fig. S1. Biosensor characterization and optimization. a** Dose-response curves of three catechol Caq101 biosensor strains using the promoters *P_PGK1_*, *P_CCW12_*, and *P_TEF1_* each containing one operator copy for reporter expression. **b** Dose-response curves of three protocatechuate PcaV biosensor strains using the promoters *P_PGK1_*, *P_CCW12_*, and *P_TEF1_* each containing one operator copy for reporter expression. **c** Dose-response curves of three protocatechuate PcaV biosensor strains using the promoters *P_PGK1_*, *P_CCW12_*, and *P_TEF1_* each containing two operator copies for reporter expression. **d** Dose-response curves of strains using the BenM^WT^ and BenM^Tisno^ biosensor variants with the *P_CYC1_* promoter containing one operator copy for reporter expression in different concentrations of ccM and adipic acid. **e** Response of biosensor candidates to 5 mM catechol, protocatechuate, ccM, and adipic acid. Datapoints are mean ± SD, *n* = 3.

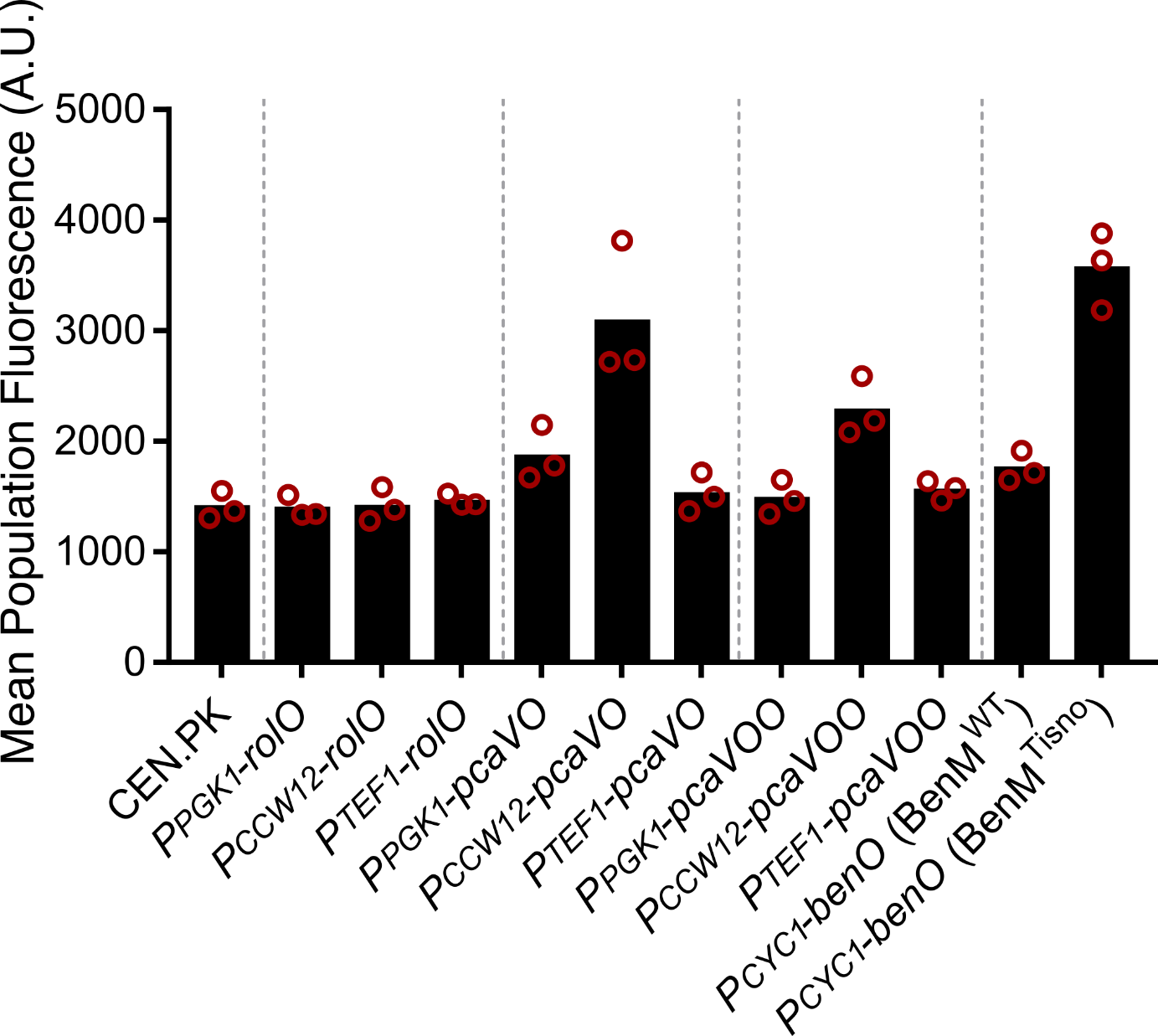

**Fig. S2. Characterization of basal fluorescence levels for all sensor strains in this work.** Mean population fluorescence of strains carrying different catechol, PCA, ccM, and adipic acid inducible promoters driving *gfp^Envy^* expression in the absence of inducers compared to a wild-type *S. cerevisiae* CEN.PK113-7D strain. Datapoints represent the mean ± SD, *n* = 3.

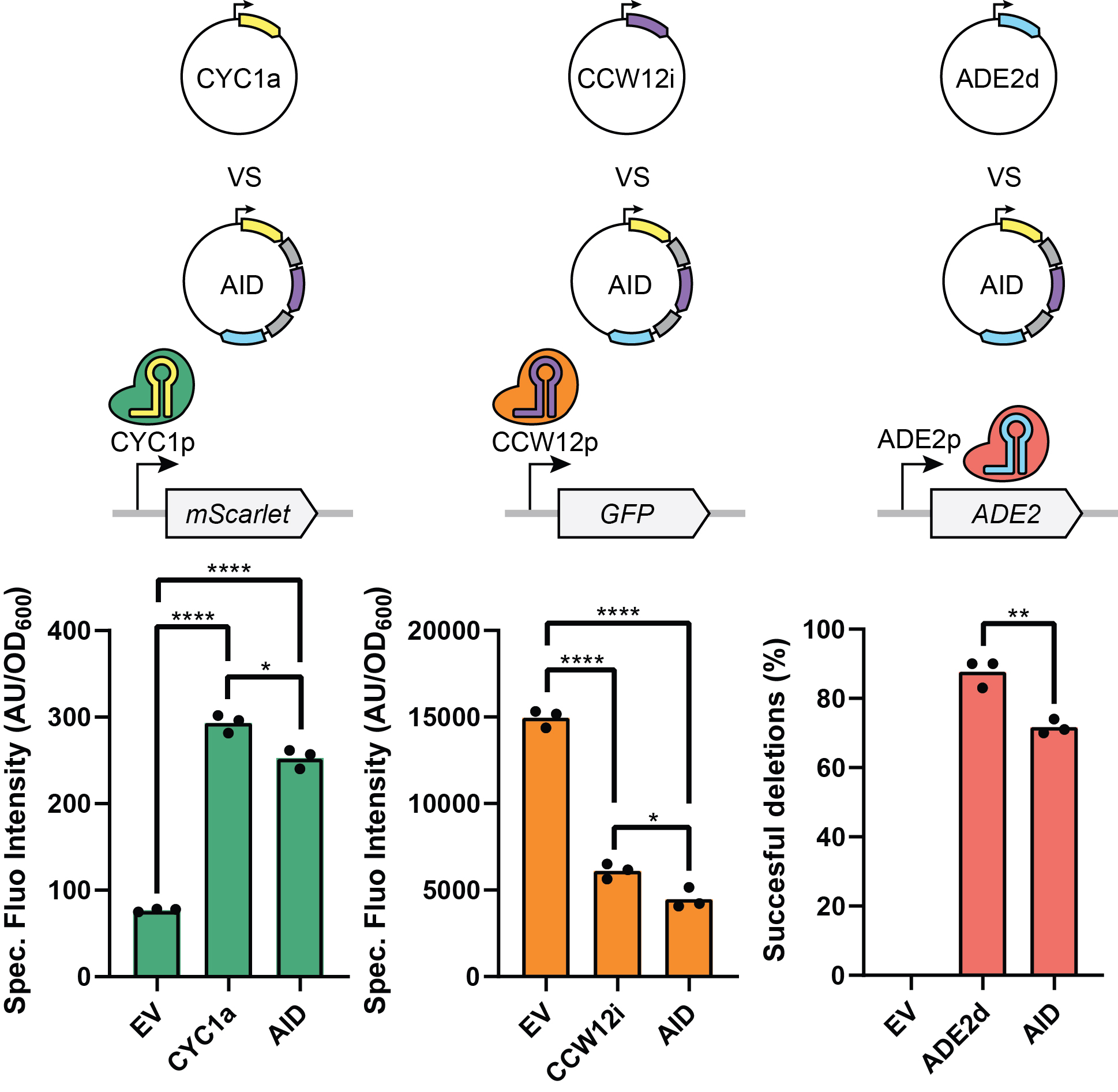

**Fig. S3. Combinatorial AID control.** Activation of mScarlet, interference on GFP, and deletion of ADE2 performed in the PCA-producing background harboring the AID Cas system (M1 strain). One single guide RNA (CYC1a, CCW12i and ADE2d) or the combinatorial system (AID) expressing the three guides simultaneously due to the tRNAgly splicing units were used to perform the different perturbations. Statistical t-tests are shown (*, *P* < 0.05; **, *P* < 0.01; ****, *P* < 0.0001).

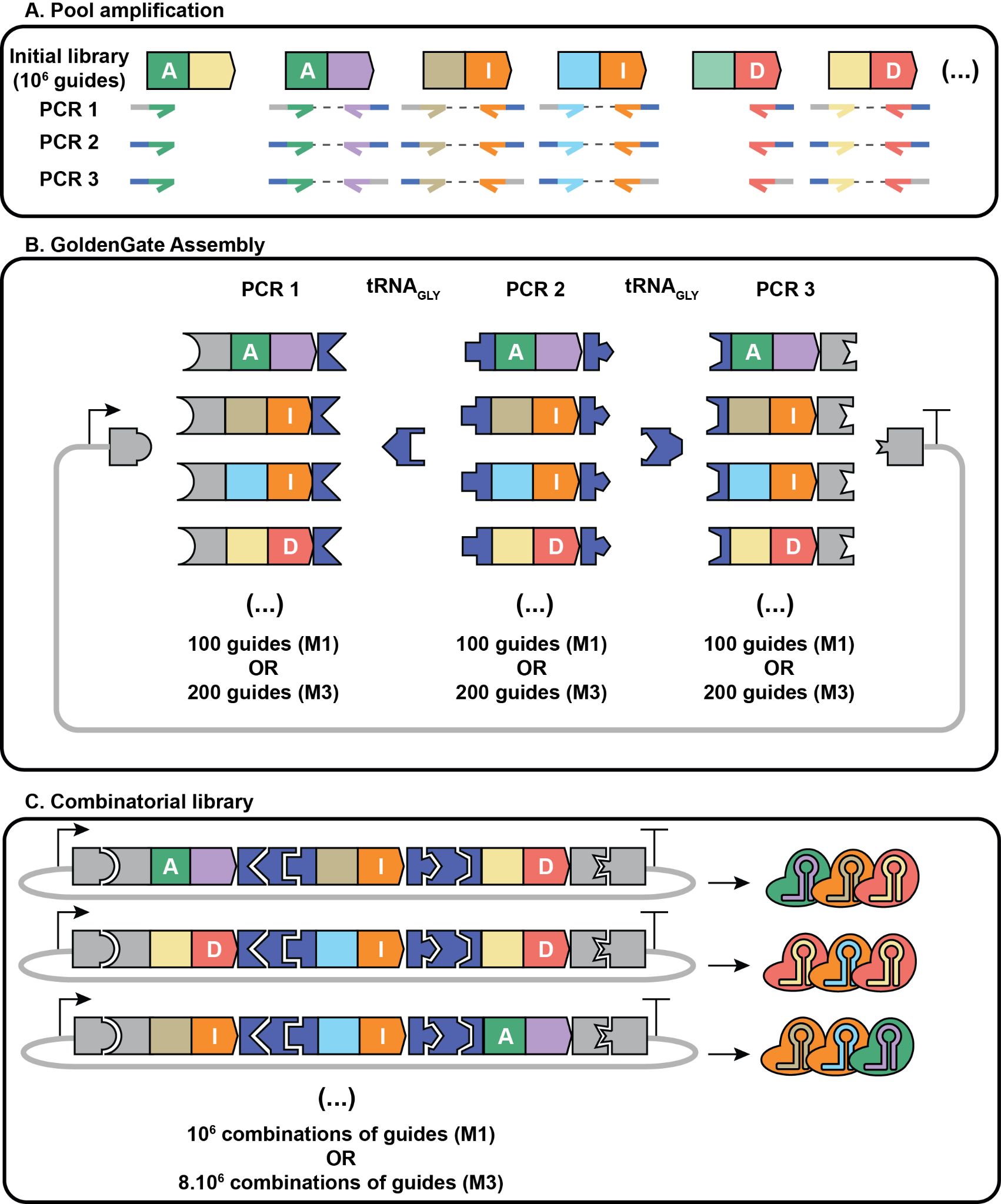

**Fig. S4. Cloning methodology for the combinatorial assembly.** The single guide RNA library used for the single-guide screen was first amplified by PCR with three primers specific to the activation, interference or deletion scaffold together with an oligo pool of primers specifically binding the guides sequences that showed previously an effect on ccM or PCA increase. Each primer was ordered in three versions, each containing a different overhang with a BsaI restriction site compatible to insert the amplicon at three different positions in the plasmid vector. After amplification, the amplicons were purified and combined together with the backbone vector and two tRNAgly parts consisting of a tRNAgly splicing unit flanked by two different BsaI overhangs. The assembly was next transformed into *E. coli* before extraction of the plasmids for yeast transformation. The combinatorial assembly made of the best 100 guides identified to boost PCA production were introduced into the M1 PCA-producing strain, and the combinatorial assembly of the 200 best guides identified to boost ccM production were introduced into the M3 ccM-producing strain.

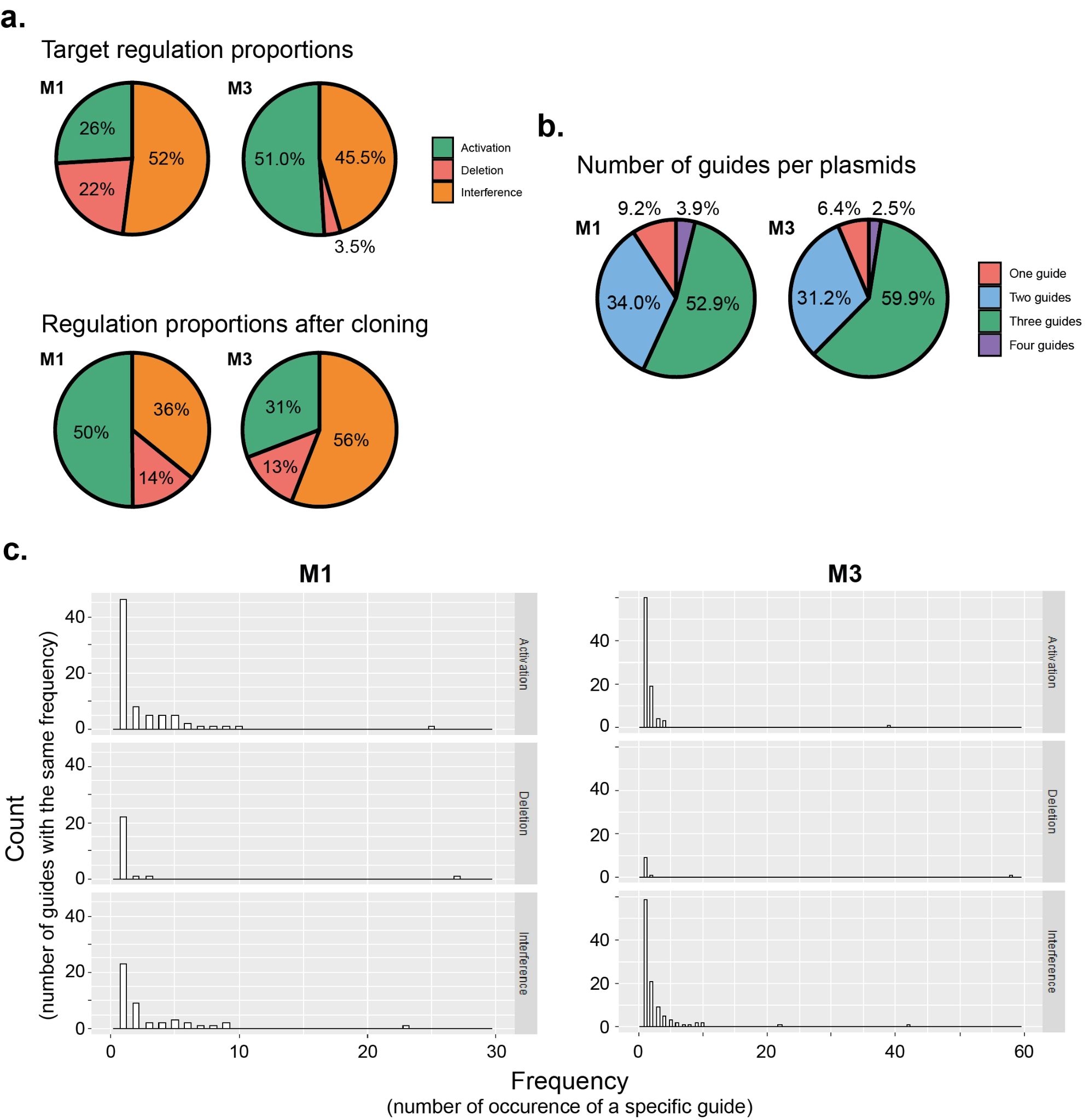

**Fig. S5. Quality control of the library.** Analysis of the sequencing of 96 random colonies from the cloned combinatorial library. **a.** Repartition of the proportions of the three different perturbations (activation, interference and deletion) in the cloned combinatorial library in comparison to the targeted repartition. The targeted repartition is defined by the 100 (M1) and 200 (M3) selected guides to build the combinatorial library. **b.** Repartition of the number of guides per plasmids. **c.** Histogram of the frequency of guides. The graphs show the number of occurrences of a specific guide (x axis), and the number of guides that have been found for each frequency.

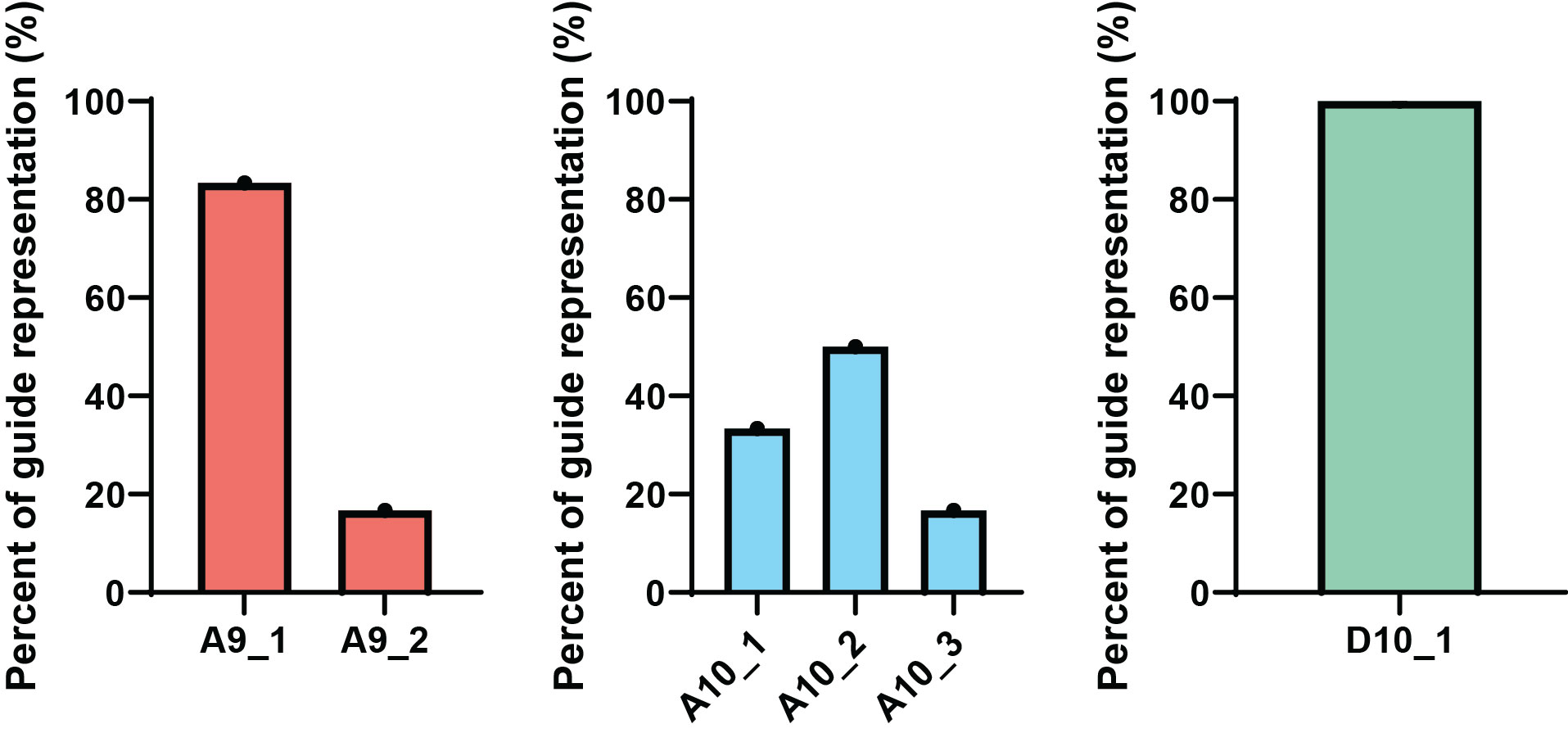

**Fig. S6. Presence of multiple gRNAs in yeast populations.** Top PCA producing colonies of yeast had their plasmids extracted. These plasmids were re-cloned into *E. coli* and sent for sequencing. Six total clones were sent for each yeast colony.

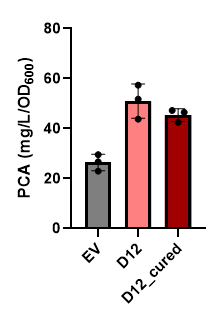

**Fig. S7. Background effect seen in Colony D12 during M1 screen.** The M1 top hit colony D12 (pink) exhibited high PCA titres compared to control (grey). However, deconvolution of the guide set did not result in any single, double, or triple guide combination increasing PCA (Fig 6). To investigate this, we cured D12 of the guide plasmid and re-analyzed production of PCA (dark red).

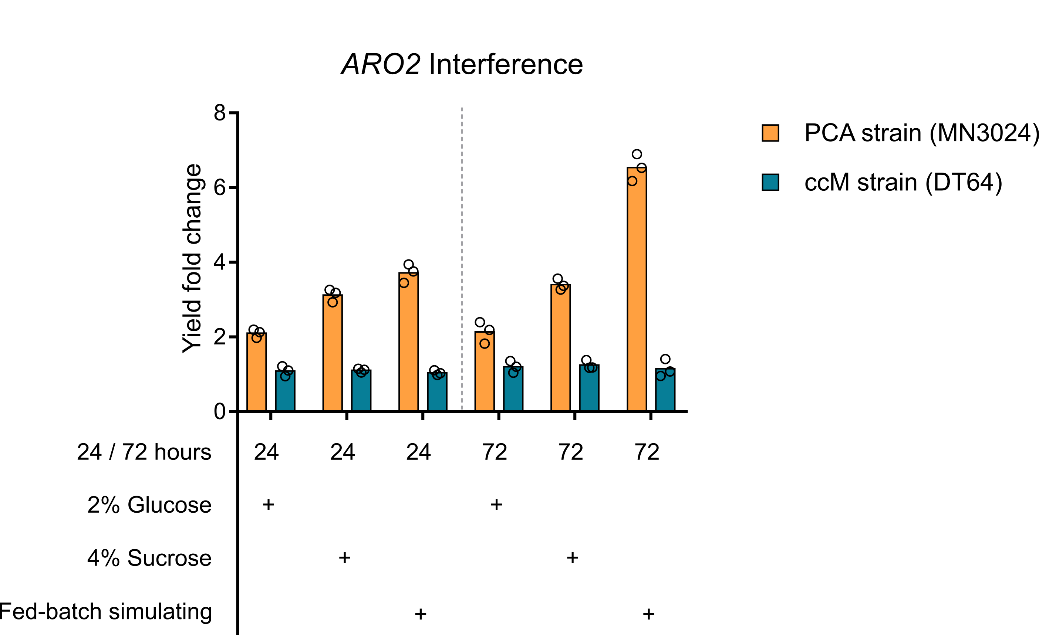

**Fig. S8. Deep-well plate bioproduction process optimization for PCA- and ccM- producing strains.** Fold-change specific yields of PCA (orange) and ccM (teal) in CRISPR-free strains harbouring an interference in expression of *ARO2* from samples taken at 24 or 72 hours using either 2% glucose, 4% sucrose, or fed-batch simulating glucose as a carbon source. Datapoints represent the mean ± SD, *n* = 3.
